# EMT-dependent cell-matrix interactions are linked to unjamming transitions in cancer spheroid invasion

**DOI:** 10.1101/2024.05.08.593120

**Authors:** Anouk van der Net, Zaid Rahman, Ankur D. Bordoloi, Iain Muntz, Peter ten Dijke, Pouyan E. Boukany, Gijsje H. Koenderink

## Abstract

The plasticity of cancer cells allows them to switch between different migration modes, promoting their invasion into the extracellular matrix (ECM) and hence increasing the risks of metastasis. Epithelial-to-mesenchymal transitions (EMT) and unjamming transitions provide two distinct pathways for cancer cells to become invasive, but it is still unclear to what extent these pathways are connected. Here we addressed this question by performing 3D spheroid invasion assays of lung adenocarcinoma (A549, epithelial) and melanoma (MV3, mesenchymal-like) cancer cell lines in collagen-based hydrogels, where we varied both the invasive character of the cells (using Transforming Growth Factor (TGF)-*β* to promote EMT and matrix metalloprotease (MMP) inhibition to block cell-mediated matrix degradation) and the porosity of the matrix. Using a quantitative image analysis method to track spheroid invasion, we discovered that the onset time of invasion mostly depended on the matrix porosity and corresponded with vimentin levels, while the subsequent spheroid expansion rate mostly depended on metalloprotease MMP1 levels and thus cell-matrix interaction. Morphological analysis revealed that spheroids displayed solid-like (non-invasive) behavior in small-pore hydrogels and switched to fluid-like (strand-based) or gas-like (disseminating cells) phases in large-pore hydrogels and when cells were more mesenchymal-like. Our findings are consistent with unjamming transitions as a function of cell motility and matrix confinement predicted in recent models for cancer invasion, but show that cell motility and matrix confinement are coupled via EMT-dependent matrix degradation.

## Introduction

The dissociation of cancer cells from the primary tumor followed by invasion through the local tumor microenvironment (TME) is a crucial first step during metastasis of solid tumors. These first stages of invasion are often achieved through epithelial-to-mesenchymal transitions (EMT) that promote cell motility^1^ and they are influenced by the composition of the TME. The TME has distinctive cellular components, including cancer-associated fibroblasts (CAFs) and immune cells, as well as non-cellular components, including cytokines, growth factors, and a unique organization of extracellular matrix (ECM)^2^. Tumor cells and CAFs remodel the ECM in the TME, generally turning the fibrous networks from healthy tissues into denser and stiffer matrices with smaller pores^3^. The resulting dysregulated ECM poses physical challenges for the invading tumor cells, as the cells have to tightly squeeze their nuclei while preventing excessive DNA damage^4, 5^.

To deal with a wide variety of dysregulated tissue organizations in the TME, tumor cells are known to adapt their phenotype and associated migration strategy in response to their local environments. Tumor cells that migrate individually can adopt three main modes of migration, which can be categorized by their dependency on cell-matrix adhesions and actomyosin contractility. Cancer cells performing mesenchymal migration pull themselves forward on matrix fibers relying on matrix adhesions and cell contractility. In low-adhesion or high-density tissues, cancer cells can migrate through either amoeboid or lobopodial migration and increase their deformability^5^. Next to these individual cell motility modes, cancer cells can also switch to multicellular migration modes. This collective migration strategy is linked to increased survival in the bloodstream and higher metastatic potential^6^. The phenotypic plasticity of cancer cells enables switches between these different migration strategies in response to changes in ECM confinement and stiffness^7, 8^, oxygen and energy deprivation^9^ and cytokines^10^.

While the role of intracellular signaling pathways (such as responses to TGF-*β*, epidermal growth factor, or cytokines released from CAFs) and the expression of EMT markers^11, 12^ in cancer cell invasion are relatively well studied, the molecular mechanisms behind migration mode switches in response to different TMEs remain poorly understood^13^. The increasing attention for the biophysical aspects of tumor invasion in recent years has revealed that matrix confinement, determined by ECM protein density and porosity, is an important regulator of cell migration mode switches^7, 14, 15^. Mechanistically, the link between ECM confinement and cancer cell invasion has been explained by describing tumors as active liquid crystalline materials that undergo solid-to-liquid-like transitions during jamming conversions^5, 16^ that are determined by cell-cell adhesions and ECM confinement^17^. Lattice-gas cellular automaton simulations suggested a theoretical jamming phase diagram that describes how variations in both cell-cell adhesions and ECM density (and hence confinement) determine unjamming transitions from a state where no invasion occurs (solid-like phase) towards collective cell invasion (liquid-like phase) or individual cell invasion (gas-like phase)^17^. Later agent-based simulations predicted a non-equilibrium phase diagram that describes these unjamming transitions in terms of ‘cell motility‘ and ‘matrix density‘^18^.

While this conceptual framework is appealing, it is complicated by the fact that these two variables, cell motility and ECM confinement, are not independent. They are coupled since cancer cells can actively modify the ECM composition and architecture, while conversely the ECM can induce changes in cell motility^19^. Cancer cells adhere to the ECM via integrin receptors, activating intracellular signalling pathways that trigger cellular responses such as ECM deposition and the release of matrix metalloproteases (MMPs)^17, 20^. MMPs are proteolytic enzymes secreted by cancer cells to break down ECM proteins such as collagen, forming proteolytic tracks that facilitate cancer cell invasion^21^. In addition, upregulation of collagen deposition by tumor cells (and CAFs) can also promote cell invasion with the formation of specialized anisotropic migration tracks^22, 23^. These complex events together contribute to an abnormal ECM composition, structure and stiffness, which in turn influences cell invasion^2, 24^. In addition, these changes in the ECM alter other cellular characteristics such as cell proliferation, differentiation and cell mechanics, all of which also impact cancer initiation and progression^25–28^. Changes in ECM stiffness and porosity can affect the mode of migration of cancer cells^21, 29^ and their invasion capacity^27, 30–33^. Moreover, the ECM can also affect cell migration through the storage and release of growth factors^34^, such as the latent form of transforming growth factor-*β* (TGF-*β*)^35–37^. TGF-*β* is a well known cytokine found in the TME that triggers the upregulation of mesenchymal-associated protein markers such as vimentin, and the down-regulation of epithelial protein markers such as E-cadherin, promoting cell motility and invasion through EMT^38, 39^. Furthermore, TGF-*β* also stimulates the release of MMPs from cancer cells, promoting the formation of proteolytic tracks that promote invasion^21^. The role of TGF-*β* in tumor invasion is a clear example of the dynamic reciprocity of cell-ECM interactions during invasion.

Although much work has been done on elucidating the different matrix properties that influence cancer cell invasion, the reported correlations can be inconsistent. For example, increased matrix stiffness correlates with enhanced invasion in 3D *in vitro* models of epithelial cancers in 64% of the studies, but it is also negatively correlated in 36% of the studies^40^. This variability is probably due to the diversity in experimental set-ups. Especially the time scale over which invasion is monitored is likely important, since invasion is a highly dynamic process. Because of these inconsistencies, there are still many open questions about the roles of matrix properties in invasion, especially in combination with reciprocal cell-ECM interactions where cell motility and matrix properties change in concert. To gain a better understanding of how cell-matrix interactions control cell invasion, we need cellular *in vitro* model systems that decouple interconnected matrix properties such as stiffness and porosity^41^, differentiate between different stages of invasion, and analyse how matrix remodelling and cell signalling impact unjamming phase transitions during invasion.

In this study, we aimed to understand the effect of mechanochemical coupling between matrix confinement and cell motility on tumor invasion. For this, we analysed the invasion efficiencies and migration modes of melanoma and lung cancer cells in 3D spheroid-hydrogel assays. We related these phenotypic characteristics to protein levels of EMT markers (vimentin and E-cadherin) and MMPs and to the unjamming transitions that describe invasion behaviour^18^. To tune matrix confinement over a wide range, we used two types of hydrogels, natural collagen (bovine type I) and semi-synthetic gelatin methacrylate (GelMA), at different concentrations. GelMA matrices contained pore sizes that were substantially smaller (≃ 10 nm range) than average cell bodies (≃ 10 micrometer range) so cell motility requires matrix degradation, while the pore sizes of collagen type I matrices (≃ micrometric) allowed for cell motility through a combination of cell squeezing and matrix degradation. We systematically compared the invasive capacity of epithelial-like cancer cells (A549, lung adenocarcinoma) and mesenchymal-like, highly metastatic cells (MV3 melanoma). Furthermore, we explored how cell-mediated matrix degradation influences invasion trends, by studying cancer cell invasion upon TGF-*β* or MMP inhibitor treatments. Using a new image analysis method to analyze the spatiotemporal characteristic of spheroid invasion, we demonstrate that the onset of spheroid invasion is regulated by initial matrix confinement and vimentin, while the rate of invasion and unjamming transitions correspond with MMP1 levels. Our findings reveal that EMT-regulated cell-ECM mechanochemical interactions play an important role in 3D spheroid unjamming transitions.

## Results

### Design and characterization of the 3D spheroid invasion model

To identify the impact of ECM confinement on the invasion of cancer cells with different metastatic potentials, we performed 3D spheroid invasion assays using two different human cancer cell lines and two different types of hydrogels. MV3 (melanoma) cells were chosen as they are highly metastatic and mesenchymal-like, in contrast to the epithelial character of A549 (lung carcinoma) cells (see schematic in Fig. 1A). We confirmed the mesenchymal and epithelial features of the two cell lines by bright-field imaging of the cell morphology (see Fig. S1A) and Western blot analysis of the protein levels of E-cadherin and vimentin (see Fig. S1B,C). For hydrogel systems, we chose collagen type I and GelMA hydrogels, because of their contrasting fibrous open-mesh network structure versus dense confining microenvironment, respectively (Fig. 1Bi). Two different concentrations for each type of hydrogel were characterized for stiffness and for the pore size and corresponding fiber density. Measurements of the storage modulus (G’) from small amplitude oscillatory shear rheology showed that the hydrogels covered a wide range of stiffnesses from 10^1^ Pa to 10^3^ Pa (see Fig. 1C). Pore size measurements were conducted through analysis of confocal microscopy reflection images for collagen networks (Fig. 1Bii,iii and Fig. S2) and through permeability measurements for GelMA hydrogels (Fig. 1Biv,v and Table S1). The GelMA hydrogels had pore sizes in the ≃ 10 nanometer range, two orders of magnitude smaller than the micrometric pores of collagen hydrogels (Fig. 1D). Furthermore, for both hydrogel types, the pore size decreased with increasing biopolymer concentration (30 mg/mL versus 50 mg/mL for GelMA; 2.4 mg/mL versus 8 mg/ml for collagen). As an independent test of matrix porosity, we also compared the fiber densities of the four hydrogels by measuring the void fraction in confocal reflection microscopy images. These estimations showed that the hydrogels with the smaller pores indeed had the highest matrix fiber densities (see Fig. S3). These results demonstrate that the ECM-mimicking hydrogels in this study offer a wide range of levels of cell confinement, from full confinement of cells in GelMA hydrogels to partial confinement of cells in the collagen hydrogels.

**Figure 1.**
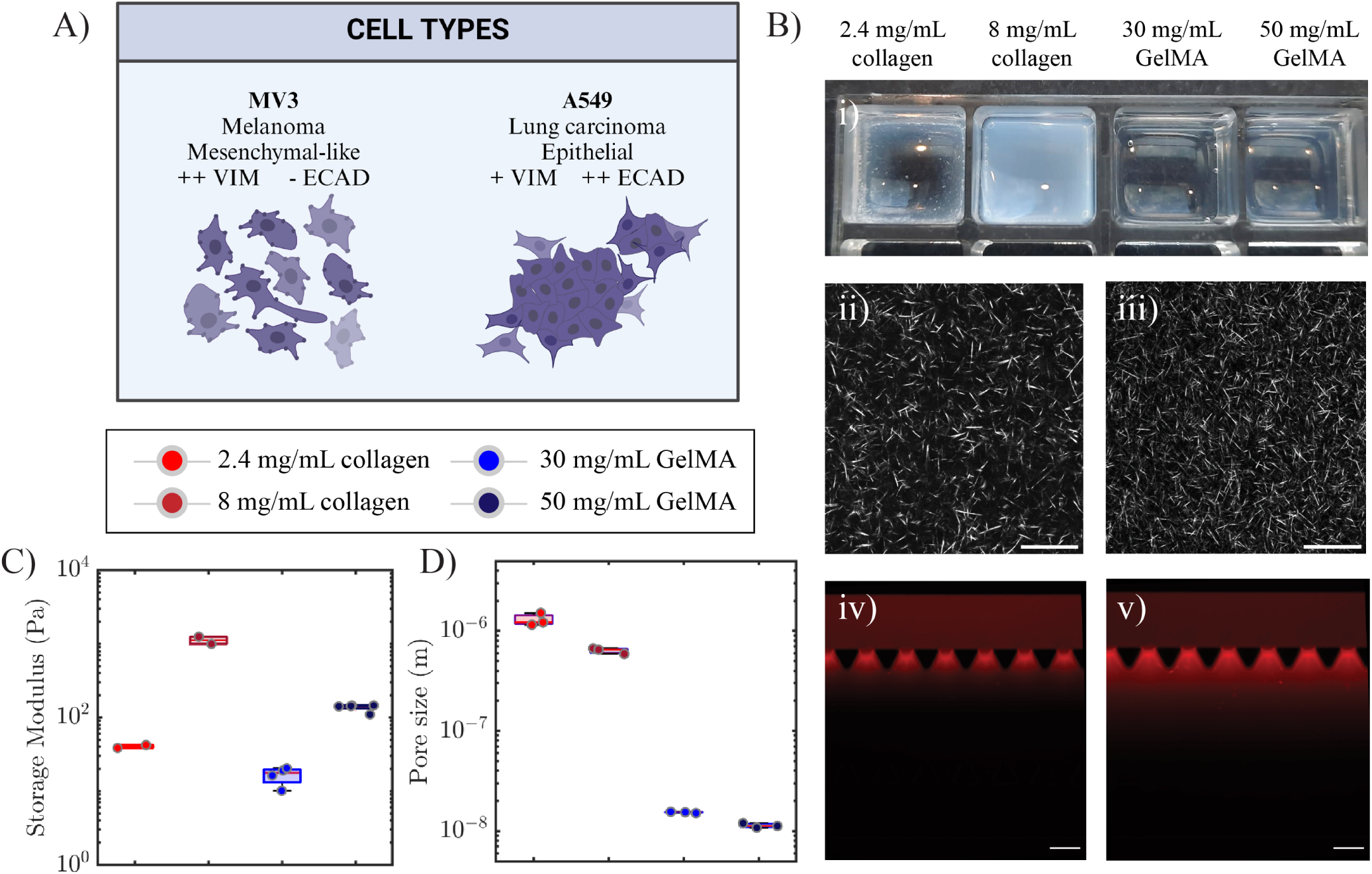
Characterization of the 3D spheroid invasion model. (A) Schematic showing the distinctive features of MV3 melanoma cells and A549 lung carcinoma cells. MV3 cells are mesenchymal-like, with high vimentin expression (++VIM) and low E-cadherin expression (-ECAD), whereas A549 cells are epithelial-like, with lower vimentin (+VIM) and high E-cadherin (++ECAD) expression and more cell-cell interactions. (B) Structural characterization of the ECM-mimicking hydrogels. (i) Photograph of the four hydrogels in an ibidi 8-well slide (without spheroids), scale bar = 10 mm. The collagen hydrogels are more turbid than the GelMA hydrogels because of the larger width of collagen fibers as compared to GelMA strands. Confocal reflection images of collagen at concentrations of (ii) 2.4 mg/mL and (iii) 8 mg/ml. Scale bars are 25 25 µm. Permeability analysis of GelMA (30 mg/mL) based on fluorescence images of Rhodamine B dye (red) at (iv) *t* = 0 and (v) *t* = 10 min, performed in a microfluidic chip, scale bar = 200 *µ*m. We inferred the hydraulic permeability from the distance travelled by the dye under a pressure gradient of 20 mbar (top to bottom) using Darcy’s law. (C) Storage shear moduli (G’) of collagen and GelMA hydrogels obtained by rheology measurements. (D) Pore sizes of collagen and GelMA hydrogels, obtained by image analysis and permeability analysis, respectively.

### Spheroid invasion depends on matrix confinement and on cell type

To understand the impact of matrix confinement (i.e., pore size) on the ability of the cells to invade, we performed invasion assays by bright-field imaging of 200 µm diameter cancer spheroids embedded in hydrogels at the spheroid equator. For MV3 spheroids in low density (2.4 mg/ml) collagen, we observed substantial cell invasion already by *t* = 8 hours (hrs), and even more invasion after 24 hrs (Fig. 2A). After 24 hrs, the spheroid was partially disintegrated, leaving an intact spheroid core surrounded by many disseminated individual cells. Therefore, the MV3 invasion assay was limited to 24 hrs. In a denser (8 mg/ml) collagen hydrogel, the MV3 spheroids showed a similar invasion pattern (Fig. S4A). In 30 mg/mL GelMA hydrogels, the MV3 spheroids also showed an isotropic invasion pattern, but invasion was delayed and the cells reached less far into the gel (Fig. 2B). In 50 mg/mL GelMA gels, we could not observe any signs of invasion on a 24 hrs time scale (Fig. S4B).

**Figure 2.**
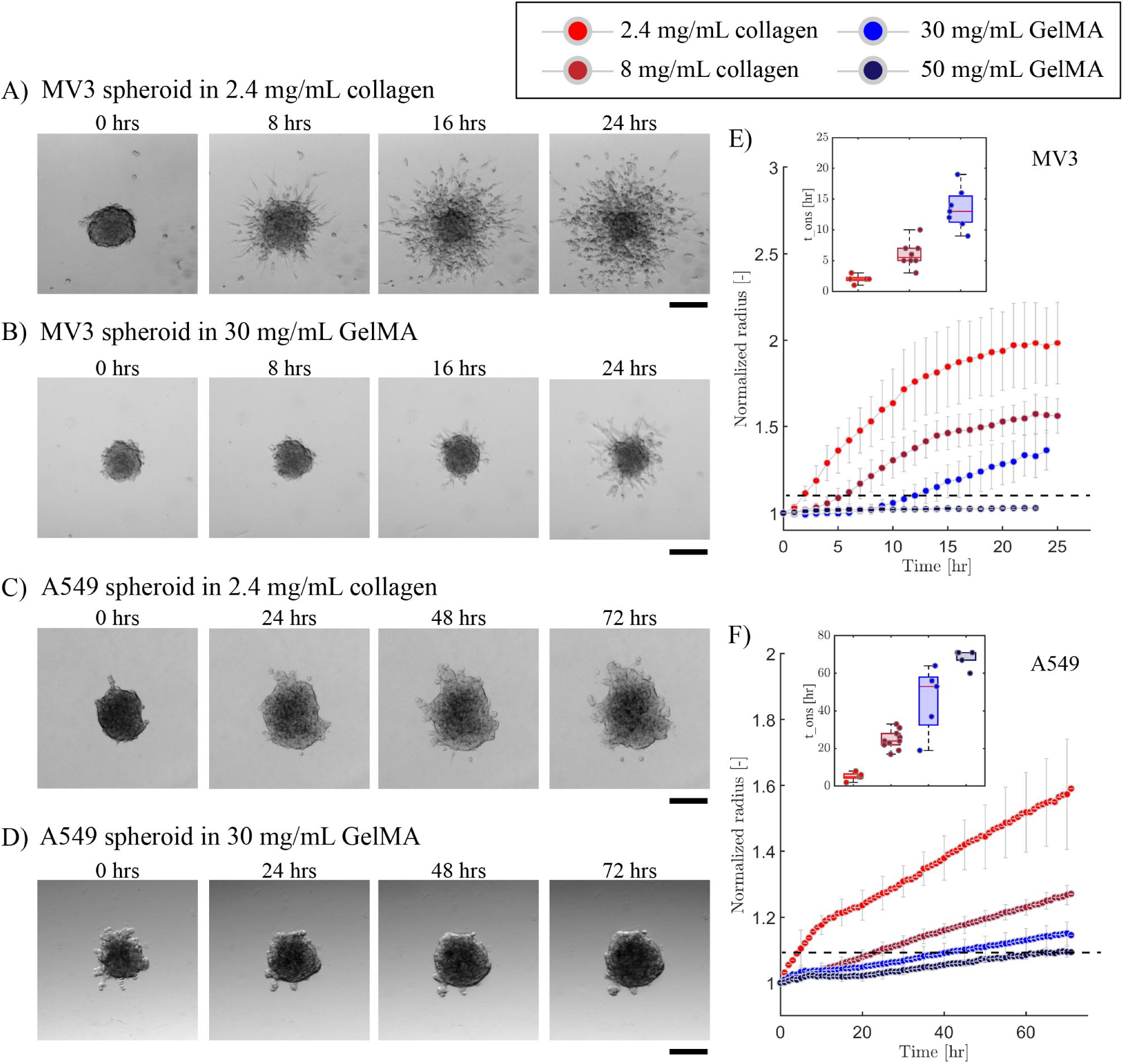
MV3 and A549 spheroid invasion in collagen and GelMA matrices. Bright-field microscopy images of spheroids at different time intervals for 24 hrs invasion of a MV3 spheroid in A) collagen (2.4 mg/mL) and (B) GelMA (30 mg/mL), and for 72 hrs invasion of an A549 spheroid in (C) collagen (2.4 mg/mL) and (D) GelMA (30 mg/mL). Scale bars: 200*µ*m. Time evolution of the normalized effective circular radius of (E) MV3 and (F) A549 spheroids, for different matrix compositions (see legend on top). Horizontal black dashed lines indicate the onset time for invasion, defined as the time where the normalized radius reaches a value of 1.1. Data are averages with standard deviations of N = 4-10 spheroids (from n = 2 independent experiments). Insets in (E) and (F) show the onset time (*t_ons_*) of invasion in the different matrices (same color code as main plots).

The A549 spheroids showed a very different invasion behavior as compared to MV3 spheroids. Even in the least confining hydrogel (2.4 mg/mL collagen), the A549 spheroids remained compact and developed only a small number of multicellular protrusions from sites that were non-uniformly distributed around the spheroid periphery. Also, there were much fewer disseminated cells (Fig.2C). The A549 spheroids furthermore showed a much longer onset time for invasion (*t ≈* 24 hrs) compared to the MV3 spheroids (t *≈* 8 hrs) in the same matrix. To allow sufficient time for cell invasion, we therefore performed 72 hrs of time-series imaging for A549 spheroids. In denser 8 mg/mL collagen hydrogels and GelMA hydrogels (30 mg/mL and 50 mg/mL), A549 spheroids showed hardly any protrusions nor disseminated cells even after 72 hrs (Fig.2D for 30 mg/mL GelMA and Fig. S5 for 8 mg/mL collagen and 50 mg/mL GelMA). These findings demonstrate that the mesenchymal-like MV3 cells are more invasive than the epithelial-like A549 cells and that invasion in both cases is impacted by the matrix density.

To quantify the effects of cell type and matrix density on spheroid invasion, we tracked the spheroid periphery for each time point through automated image analysis of the bright-field time-lapse images. To account for varying spheroid shapes and initial sizes, we computed the effective circular radius normalized by its initial value at *t* = 0 hr. For the MV3 spheroids, we found a clear trend of decreasing cell invasion with decreasing matrix pore size, going from 2.4 mg/mL collagen to 8 mg/mL collagen and to 30 mg/mL GelMA and finally to 50 mg/mL (Fig. 2E). In the densest hydrogel (50 mg/mL GelMA), no appreciable invasion took place. The A549 spheroids showed the same trend as MV3 spheroids as a function of matrix pore size and likewise showed complete inhibition of invasion in 50 mg/mL GelMA (Fig. 2F). These invasion trends strongly suggest that the matrix pore size is an important determinant of the spheroid invasion capacity. By contrast, the matrix stiffness appeared to have little predictive value, since collagen (2.4 mg/mL) gels and GelMA (30 mg/mL) gels had very different effects on spheroid invasion despite their comparable elastic modulus (40 Pa for collagen (2.4 mg/mL) and 15 Pa for GelMA (30 mg/mL). It is also interesting to note that the GelMA (50 mg/mL) gel had a rather low stiffness of just G’ = 200 Pa, yet showed complete inhibition of invasion for both cancer cell types.

### Increased matrix confinement delays the onset of cell invasion

The invasion time curves clearly showed a delay time before measurable spheroid invasion occurred, followed by a steady increase of normalized spheroid radius (Fig. S6). To quantify this delay, we defined the onset time for invasion (*t_ons_*) as the time point at which spheroids achieved a 10% increase in normalized radius, shown by the horizontal dashed lines in Fig. 2E,F. For MV3 spheroids in collagen, the delay time was shortest (*t_ons_*= 3 hrs) in 2.4 mg/mL collagen, somewhat longer (*t_ons_* = 5 hrs) in 8 mg/mL collagen, and even longer (*t_ons_* = 11.5 hrs) in 30 mg/mL GelMA (inset of Fig.2E). For A549 spheroids, the onset times were always longer than for MV3 spheroids, but there was a similar trend of increasing onset time with decreasing pore size, from *t_ons_*) = 15 hrs for 2.4 mg/mL collagen, to *t_ons_* = 24 hrs for 8 mg/mL collagen, and *t_ons_*= 57 hrs for 30 mg/mL GelMA (inset of Fig.2F). In 50 mg/mL GelMA, neither the MV3 nor the A549 spheroids ever reached the threshold of a 10% increase in normalized spheroid radius. Based on these results, we conclude that the matrix porosity affects the invasion capacity of the spheroids by controlling the onset time of invasion.

### MMP-mediated matrix degradation promotes spheroid invasion

Since matrix porosity strongly impacted the invasion capacity of the spheroids, we hypothesized that MMP secretion might be an important determinant of the invasion capacity. To test this hypothesis, we decided to tune the ability of the cells to degrade the matrix by using treatments with TGF-*β* to promote MMP secretion^42, 43^. Bright field imaging showed that TGF-*β* strongly promoted invasion of MV3 spheroids in 8 mg/mL collagen (Fig. 3A, left). After 24 hrs, the spheroid was disintegrated, with a partially diminished spheroid core. An even more drastic effect was seen for MV3 spheroids in 2.4 mg/mL collagen, where TGF-*β* treatment resulted in spheroid disintegration already after 10 hrs and caused the spheroid core to sink to the bottom of the well (Fig. S7A). For MV3 spheroids in GelMA (30 mg/mL), we observed a less drastic effect of TGF-*β* addition (Fig. 3A, right). After 24 hrs, the spheroid core did not disintegrate, but we did observe invasion and the presence of individual disseminated cells. For A549 spheroids, we observed qualitatively similar effects of TGF-*β* treatment. In 2.4 mg/mL collagen, spheroid invasion now was already substantial after 24 hrs, with a completely disintegrated spheroid core that sunk to the bottom of the well (Fig. S7B). In denser (8 mg/mL) collagen and in GelMA (30 mg/mL), the A549 spheroids displayed protrusions into the matrix but without any individual cell dissemination (Fig.3B). For both cell types, TGF-*β* treatment was ineffective to produce invasion in 50 mg/mL GelMA hydrogels (Fig. S8). These findings are consistent with our hypothesis that MMP-mediated degradation can promote spheroid invasion in dense matrices, but only up to a point.

**Figure 3.**
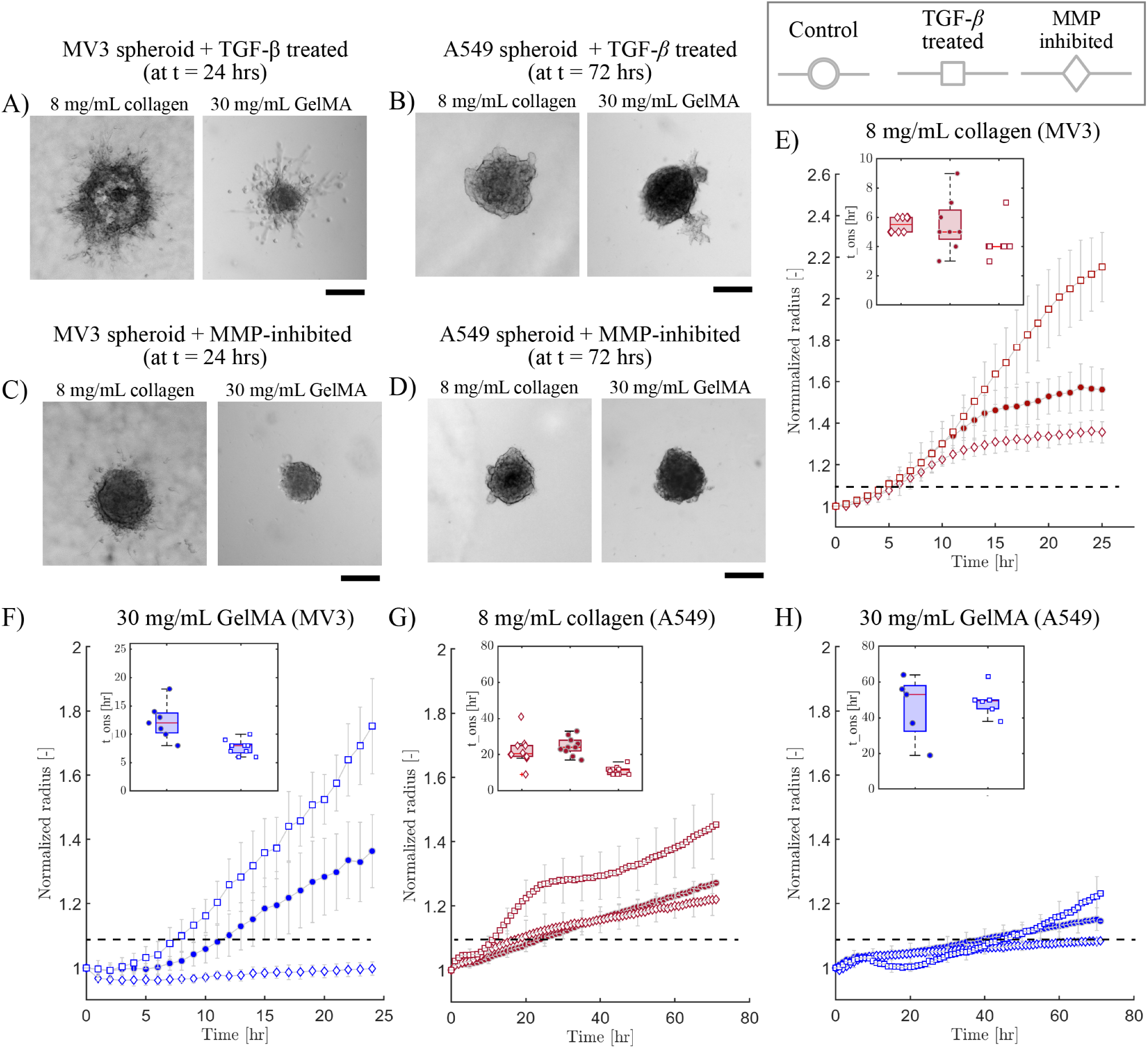
Spheroid invasion in different matrices in the presence of either TGF-*β* or the broad-spectrum MMP-inhibitor Batimastat. Bright field images of spheroids after invasion (24 hr for MV3 spheroids, 72 hrs for A549 spheroids) for TGF-*β* -treated (A) MV3 spheroids and (B) A549 spheroids, and Batimastat-treated (C) MV3 spheroids and (D) A549 spheroids. In all cases, data are shown for spheroids in 8 mg/mL collagen (left) and in 30 mg/mL GelMA (right). Scale bars are 200 *µ*m. Time evolution of normalized effective circular radius of the spheroids for (E) MV3 spheroids in collagen (8 mg/mL), (F) MV3 spheroids in GelMA (30 mg/mL), (G) A549 spheroids in collagen (8 mg/mL) and (H) A549 spheroids in GelMA (30 mg/mL). Horizontal black dashed lines indicate the onset time for invasion, defined as the time where the normalized radius reaches a value of 1.1. Data are averages with standard deviations of N = 5-10 spheroids in 2 independent experiments. Insets: invasion onset times (*t_ons_*).

Conversely, to test the effect of reduced MMP-mediated matrix remodelling, we repeated the spheroid invasion assays in the presence of the broad-spectrum MMP-inhibitor Batimastat to inhibit MMP activity^44, 45^. Bright field imaging showed a subtle effect on MV3 spheroids in 8 mg/mL collagen, where MMP-inhibitor treatment resulted in shorter cellular protrusions and fewer disseminated individual cells as compared to control conditions (Fig. 3C left). A similar behavior was seen for MV3 spheroids in 2.4 mg/mL collagen (see Fig. S7A). The effect of the MMP-inhibitor was more pronounced for MV3 spheroids in GelMA (30 mg/ml), where invasion was completely inhibited over the 24 hr time window of observation (Fig. 3C right). For the A549 spheroids, MMP-inhibitor treatment completely suppressed spheroid protrusions and cell dissemination, both in collagen hydrogels (Fig. 3D left and Fig. S7B) and in GelMA hydrogels (Fig. 3D right).

To quantify the observed effects of TGF-*β* and MMP-inhibitor treatments on invasion, we again measured the normalized spheroid radii as a function of time. Compared to control conditions, TGF-*β* treatment increased invasion of MV3 spheroids, as quantified by the final spheroid size, by 38% in collagen (8 mg/mL) and by 29% in GelMA (30 mg/mL) (Fig. 3E). In 2.4 mg/mL collagen, we observed a 20% increase in final spheroid size compared to control conditions, but we note that in this case we only quantified invasion up to 14 hours since the TGF-*β* treatment caused spheroid disintegration and sedimentation (Fig. S7C). By contrast, MMP-inhibition reduced invasion of MV3 spheroids as quantified by the final spheroid size, by 15% in 8 mg/mL collagen, 36% in 30 mg/mL GelMA, and 34% in 2.4 mg/mL collagen (Fig.S7C). The effects of the drugs were similar for A549 spheroids. TGF-*β* treatment increased invasion (after 72 hrs) by 14% in 8 mg/mL collagen (Fig. 3G) and by 10% in 30 mg/mL GelMA (Fig. 3H) compared to control conditions. By contrast, MMP-inhibition reduced invasion by 4% in 8 mg/mL collagen, 7% in 30 mg/mL GelMA (Fig. 3G, H), and 38% in 2.4 mg/mL collagen (Fig. S7D). These findings show that MMP-mediated degradation promotes spheroid invasion in dense matrices (for both cell types) and that it is absolutely required for invasion in dense GelMA gels that present pores smaller than the cell size.

### MMP-mediated remodelling has distinct effects on invasion onset time versus expansion rate

#### MMP-mediated remodelling affects the invasion onset time more strongly in GelMA than in collagen

Since we found that confinement influences invasion by delaying the onset time, we tested whether the TGF-*β* and MMP-inhibitor treatments change *t_ons_*. For MV3 spheroids in 8 mg/mL collagen, we observed a trend of reduced onset time with TGF-*β* treatment (*t_ons_*= 4 hrs) and increased onset time with MMP inhibition (*t_ons_* = 5.5 hrs) compared to the control condition (*t_ons_* = 5 hrs), but the differences were not statistically significant (inset of Fig. 3E). For MV3 spheroids in 2.4 mg/mL collagen, we similarly observed a slight increase in onset time (*t_ons_* = 3 hrs) with MMP inhibition compared to control and TGF-*β* treated spheroids (both *t_ons_* = 2 hrs) (insets of Fig. S7C, D). For MV3 spheroids in 30 mg/mL GelMA, TGF-*β* treatment significantly reduced the onset time (*t_ons_*= 8 hrs) compared to control conditions (*t_ons_* = 11.5 hrs), while MMP-inhibition resulted in complete inhibition of invasion (inset of Fig. 3F). Thus, MV3 spheroid invasion into this dense matrix was dependent upon MMP-mediated degradation. In 50 mg/mL GelMA the MV3 spheroids never invaded, even when TGF-*β* was added to upregulate MMP expression (Fig. S8C).

For A549 spheroids in 8 mg/mL collagen, we observed a stronger effect of TGF-*β* treatment on the invasion onset time (*t_ons_* = 11.5 hrs) compared to control conditions (*t_ons_* = 24 hrs) than for MV3 spheroids (inset of Fig. 3G). However, MMP-inhibition did not significantly change the onset time (*t_ons_* = 21 hrs). In 2.4 mg/mL collagen, MMP-inhibition did have a major influence on the onset time, increasing it from *t_ons_* = 5 hrs to *t_ons_* = 30 hrs (Fig. S7D). For A549 spheroids in 30 mg/mL GelMA, we did not see a significant difference between the onset time with TGF-*β* (*t_ons_* = 53 hrs) versus control conditions (*t_ons_* = 47 hrs), but MMP-inhibition completely suppressed invasion (inset of Fig. 3H). This finding again shows that invasion into the dense GelMA networks requires MMP-mediated remodelling. In 50 mg/mL GelMA, TGF-*β* -treatment caused a slight (2%) increase in final normalized spheroid radius compared to control and MMP-inhibited spheroids (Fig. S8D) and a small reduction of the onset time (*t_ons_* = 57 hrs) compared to control and MMP-inhibited conditions (*t_ons_* = 67.5 hrs and 68 hrs, respectively, inset of Fig. S8D).

### MMP-mediated remodelling impacts spheroid expansion rates

The TGF-*β* and MMP-inhibitor treatments strongly affected the final spheroid radii yet only minimally affected the invasion onset time, particularly in collagen. To understand this discrepancy, we analyzed the rate of expansion of the spheroids after the onset of invasion. We determined the expansion rates as the power-law slopes of the normalized radius versus time curves. Note that we did not perform any expansion rate analysis for MV3 and A549 spheroids in 50 mg/mL GelMA, since invasion was negligible under these conditions. For MV3 spheroids in collagen, the spheroid radius reached a plateau value around *t* = 20 hrs (Fig. 4), likely due to dissociation of cells from the spheroid that are not included in the normalized spheroid radius (Fig. S9). We therefore determined the expansion rate over an intermediate time range between the onset time and the time where the normalized radius saturated (Fig. S6). For A549 spheroids, the equivalent circular radius showed a steady increase until 72 hrs since there was little to no dissociation of cells, so we calculated the expansion rate from the onset of invasion till the end time-point (Fig. S10). The spheroid expansion rates measured under all the different conditions are summarized in Table 1.

**Figure 4.**
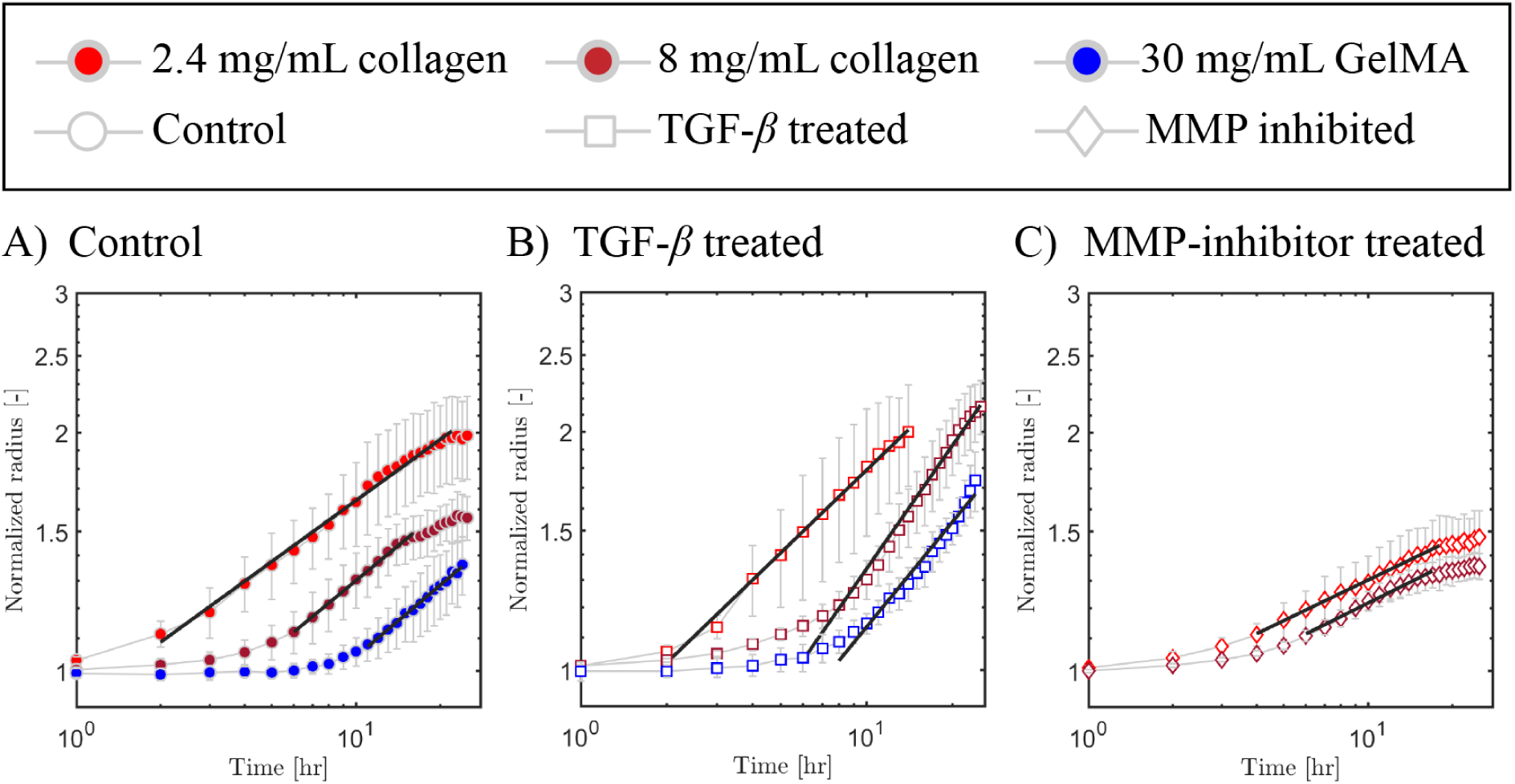
Expansion rates of MV3 spheroids. We determined the expansion rates by fitting the increase in normalized spheroid radius with time (symbols) to a power law (black lines) over the indicated temporal range. We compare MV3 spheroid invasion behavior under (A) control conditions with (B) TGF-*β* treatment and (C) MMP-inhibitor treatment. In each case, data are shown for different matrix compositions (see legend on top). Note that in panel C, we did not include MV3 spheroids in 30 mg/mL GelMA since there was no measurable invasion. Data are averages with standard deviation of N = 4-10 spheroids for each condition performed in 2 independent experiments.

**Table 1.** Expansion rates quantified as power law slope values (unit: hrs*^−^*^1^) obtained from normalized spheroid radius [-] versus time (hrs) curves for control, TGF-*β* -treated and MMP-inhibited MV3 and A549 spheroids in different hydrogel compositions. The table represents the average slope value for spheroids in collagen (2.4 and 8 mg/mL) and GelMA (30 mg/mL) with upper and lower limit of error (in square brackets). NA (not applicable) means no slope analysis was performed since no invasion was observed (MMP-inhibited MV3 and A549 spheroids in 30 mg/mL GelMA).

| MV3 spheroid expansion rates |  |  |  |
| --- | --- | --- | --- |
|  | 2.4 mg/mL collagen | 8 mg/mL collagen | 30 mg/mL GelMA |
| Control | 0.26 [0.24 – 0.27] | 0.29 [0.28 – 0.31] | 0.29 [0.28 – 0.31] |
| TGF-β treated | 0.34 [0.33 – 0.36] | 0.51 [0.49 – 0.55] | 0.44 [0.40 – 0.48] |
| MMP-inhibited | 0.17 [0.16 – 0.18] | 0.17 [0.16 – 0.19] | NA |
| A549 spheroid expansion rates |  |  |  |
| Control | 0.15 [0.14 – 0.16] | 0.14 [0.13 – 0.14] | 0.1 [0.1 – 0.2] |
| TGF-β treated | 0.37 [0.35 – 0.39] | 0.19 [0.19 – 0.2] | 0.23 [0.21 – 0.25] |
| MMP-inhibited | 0.07 [0.07 – 0.08] | 0.08 [0.07 – 0.08] | NA |

The expansion rate of the MV3 spheroids was enhanced by TGF-*β* treatment (Fig. 4B) and reduced by MMP-inhibition (Fig. 4C) as compared to control conditions (Fig. 4A). The expansion rate of the A549 spheroids was similarly impacted by TGF-*β* and MMP-inhibitor treatments. Strikingly, for both cell types, the spheroid expansion rate in control conditions was independent of matrix composition, showing comparable values in collagen (2.4 mg/mL and 8 mg/mL) and in GelMA (30 mg/mL). The expansion rate for MV3 spheroids was about two-fold higher than for A549 spheroids, as expected based on their more mesenchymal-like character. Upon TGF-*β* treatment, the spheroid expansion rates significantly increased for both cell types, by about 1.5-fold for MV3 spheroids and 2-fold for A549 spheroids. The relative increase for MV3 spheroids was highest in 8 mg/mL collagen, while for A549 spheroids it was highest in 2.4 mg/mL collagen, showing a cell type-dependent factor. By contrast, MMP-inhibition caused a 1.7-fold reduction of the spheroid expansion rates for both A549 and MV3 spheroids in both collagen matrices (2.4 mg/mL and 8 mg/mL). In GelMA (30 mg/mL), there was no detectable invasion at all in MMP-inhibited conditions. Altogether, this analysis demonstrates that the expansion rate of the spheroids is strongly dependent on MMP-mediated matrix remodelling for both cell types. In collagen hydrogels, MMP-mediated matrix remodeling speeds up invasion but is not a prerequisite. However, in GelMA hydrogels, MMP-mediated matrix degradation is a prerequisite for invasion.

### TGF-***β*** and MMP inhibitor treatments affect expression of EMT markers

The MV3 and A549 spheroids showed a similar sensitivity to MMP inhibitors, but a slightly different sensitivity to TGF-*β* treatment. The response to TGF-*β* likely is a combined effect from multiple upregulated cellular processes besides enhanced MMP expression. To test the effects of the TGF-*β* as well as MMP inhibitor treatments on these phenotypic changes, we used Western blot analysis to measure the levels of MMPs and EMT marker proteins, specifically vimentin as a mesenchymal marker and E-cadherin as an epithelial marker. For both cell lines, TGF-*β* treatment resulted in significantly upregulated levels of MMP1 (Fig. 5A) and MMP2 (Fig. 5B) compared to the control samples. At the same time, TGF-*β* treatment increased vimentin expression (Fig. 5C) and reduced E-cadherin expression (Fig. 5D) in A549 cells, confirming that TGF-*β* treatment induces an EMT switch in these epithelial-like cells. For MV3 cells, TGF-*β* treatment did not affect vimentin expression (Fig. 5C) nor E-cadherin expression (Fig. 5D), confirming that these cells are intrinsically already mesenchymal-like and therefore less affected by TGF-*β* treatment than the A549 cells. The mechanism of action of TGF-*β* is through activation of the SMAD pathway^37^. To test activation of this pathway and its intrinsic activity in the two cell lines, we analyzed phosphorylated SMAD2 (pSMAD2) levels in response to 1 hr TGF-*β* treatment by Western blot analysis. We observed a clear upregulation of pSMAD2 in response to TGF-*β* treatment in both cell lines, while the SMAD2 level remained constant (see Fig. S13). This confirms that TGF-*β* treatment activated the SMAD pathway and that the intrinsic activity of the SMAD pathway in both cell lines was at a similar level that could be upregulated by TGF-*β*.

**Figure 5.**
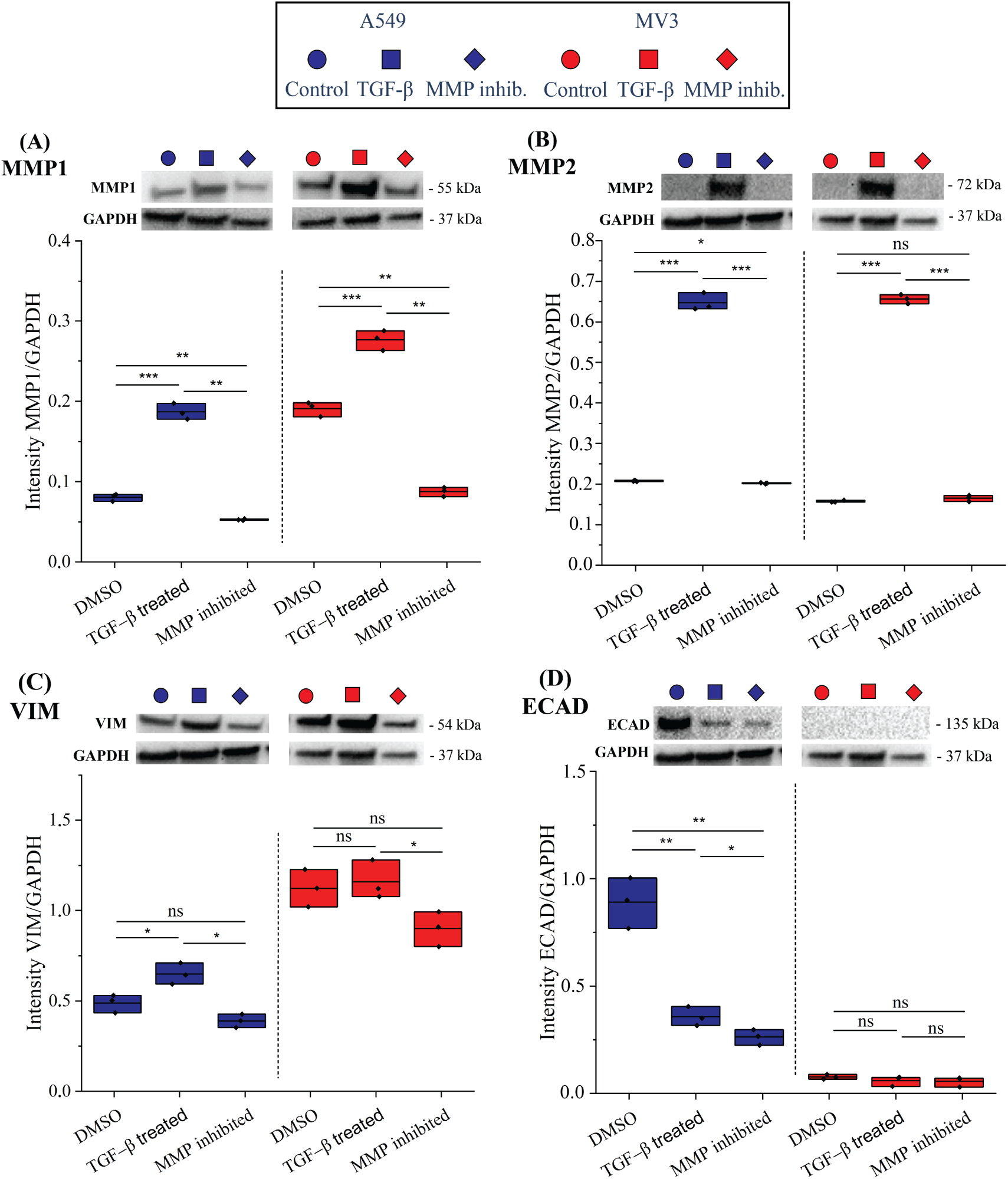
Western blot analysis of A549 and MV3 cells, showing the impact of TGF-*β* stimulation and MMP inhibition. Protein levels normalized by the GAPDH levels for A549 (blue, left side) and MV3 (red, right side) cells treated with DMSO (control conditions), TGF-*β*, or the broad spectrum MMP inhibitor Batimastat, for (A) MMP1, (B) MMP2, (C), vimentin (VIM), and (D) E-cadherin (ECAD). P-value results from t-tests are indicated by: (ns) = p*≥*0.05, (*) = p<0.05, (**) = p<0.01, (***) = p<0.001. Each condition depicts one biological sample (n=1) with three data points based on different background subtractions. The other three biological replicates (total of n=4) showed similar trends (Fig. S11A and Fig. S12A,B). The complete gels are shown in Fig. S11C,D and Fig. S12B.

Interestingly, the MMP-inhibitor also caused changes in protein expression in both cell lines. MMP inhibition reduced the expression levels of MMP1 and MMP2 (Fig. 5A,B) and decreased vimentin expression levels (Fig. 5C) in both MV3 and A549 cells compared to control conditions. MMP inhibition also reduced E-cadherin expression for A549 cells (Fig. 5D). Consistent with the reduction in E-cadherin levels in A549 cells by MMP inhibitor and TGF-*β* treatment, bright-field imaging showed that the drug-treated cells were more individual with less contact with neighboring cells compared to control conditions (see Fig. S14). In summary, the Western blot data confirm that TGF-*β* and MMP-inhibitor treatments alter MMP-levels in opposite directions and show that both treatments also impact EMT processes, especially in A549 cells.

### Invasion onset and spheroid expansion rate are correlated with matrix porosity and levels of EMT markers

The quantitative analysis of spheroid invasion showed that matrix porosity is a major determinant of the onset of cancer cell invasion. At the same time, the Western blot analysis suggests that the EMT status and MMP expression levels of the cells are also major determinants of invasion. MV3 cells were consistently more invasive than A549 cells, consistent with their higher basal levels of MMPs and the mesenchymal marker protein vimentin. TGF-*β* treatment made both MV3 and A549 cells more invasive, by upregulating MMP and vimentin expression while downregulating E-cadherin expression. Conversely, MMP inhibition made the cells less invasive by blocking MMP activity while also changing EMT marker expression. To study the interplay between matrix porosity and cell invasiveness, we prepared heat map representations to correlate the invasion onset times and the spheroid expansion rates with the cellular levels of EMT markers (*x*-axis) and with the hydrogel pore size (*y*-axis). The color code in the heat maps encodes the invasion onset time (Fig. 6A,B) and the spheroid expansion rate (Fig. 6C,D). Note that we combined the data for A549 and MV3 cells (distinguishable by symbol color) and for DMSO (control), TGF-*β* and MMP inhibitor treatments (distinguishable by symbol shape).

**Figure 6.**
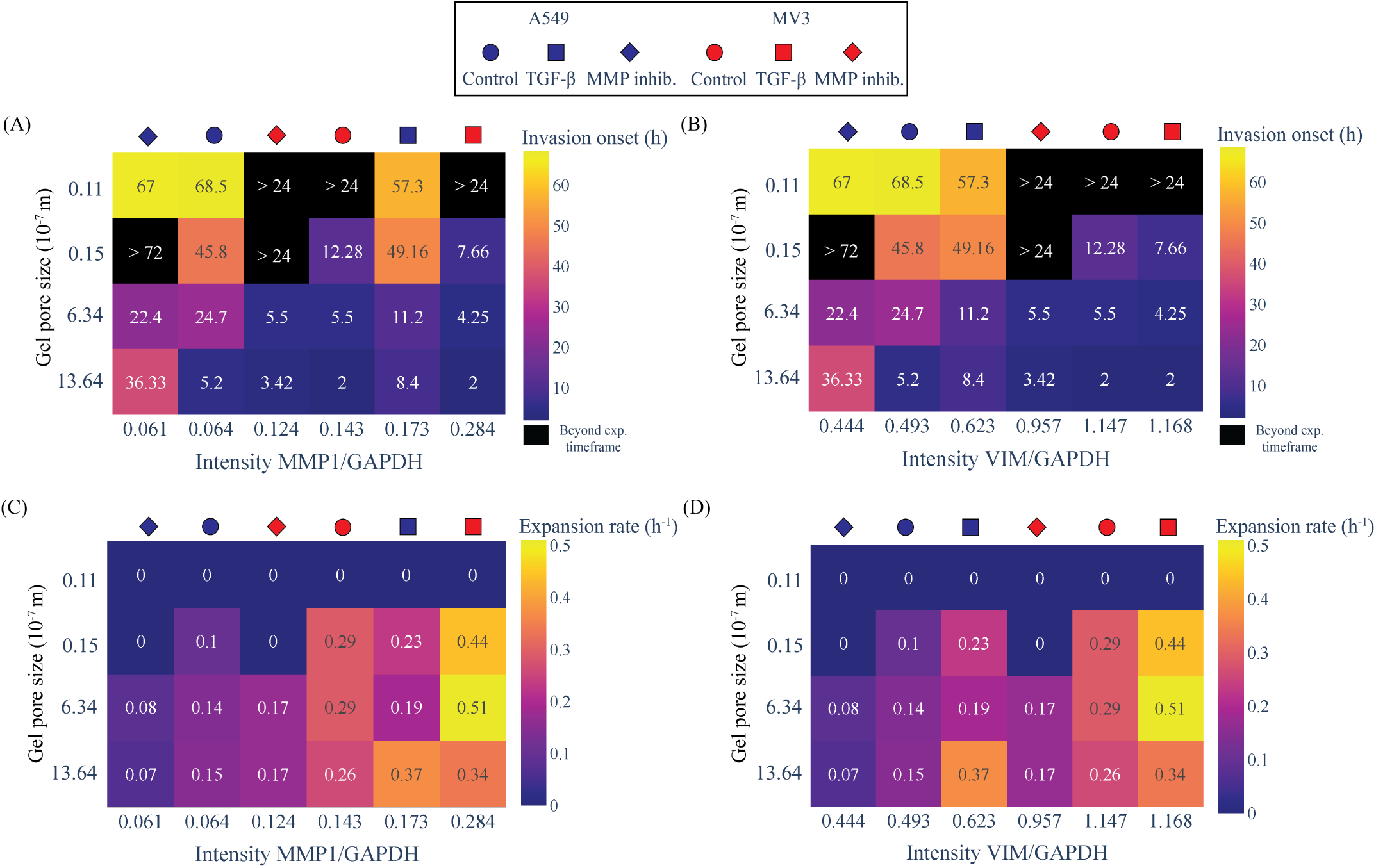
Heat map representations showing the correlation of invasion onset times and spheroid expansion rates with MMP1 and vimentin protein expression levels for MV3 and A549 spheroids. A) Dependence of the invasion onset time (indicated by the number (in units of hrs) in each square and the color code, see color bar on the right) on hydrogel pore size (*y*-axis) and expression level of MMP1 (*x*-axis). B) Corresponding heat map with the expression level of vimentin (*x*-axis). Note that spheroids that did not invade during the time frame of the assay are shown in black. C) Dependence of the spheroid expansion rate (indicated by the number (in units of hrs*^−^*^1^) in each square and the color code, see color bar on the right) on hydrogel pore size (*y*-axis) and expression level of MMP1 (*x*-axis). D) Corresponding heat map with the expression level of vimentin on the *x*-axis. Data for MV3 spheroids (red symbols above the heat maps) and A549 spheroids (blue symbols) and for different treatment conditions (see symbol shapes) were pooled. Protein expression levels were normalized by the GAPDH levels.

The heat maps reveal that the invasion onset time is mainly determined by matrix confinement, with a progressively longer delay as the pore size becomes smaller (Fig. 6A,B). In the densest 50 mg/mL GelMA hydrogels, spheroid invasion is even fully blocked under most conditions (black squares). Higher vimentin levels were correlated with shorter invasion onset times (Fig. 6B). By contrast, we did not observe any clear correlation with invasion onset time for the other tested EMT markers, MMP1 (Fig. 6A), MMP2 (Fig. S15A) and E-cadherin (Fig. S15B).

The heat maps further reveal that the spheroid expansion rate was rather insensitive to matrix confinement, except at the smallest pore sizes (50 mg/mL GelMA), where spheroid expansion was blocked (Fig. 6C,D). By contrast, the spheroid expansion rate strongly increased with increasing expression levels of MMP1 enzymes (Fig. 6C). The expansion rates did not show any clear correlation with the levels of vimentin (Fig. 6D), MMP2 (Fig. S15C) or E-cadherin (Fig. S15D). Altogether, the heat maps therefore reveal an intriguing interplay of cellular properties and matrix porosity in determining the onset of invasion and the spheroid expansion rate.

## Discussion

Here we studied how matrix porosity and cell motility parameters together determine tumor invasion by quantitative analysis of 3D spheroid invasion in collagen-based hydrogels. We found that matrix confinement and cell motility have distinct effects on the initiation of invasion and the subsequent rate of spheroid expansion. Matrix confinement and the vimentin expression level of the cells were the main determinants of the onset time of invasion. Larger pores and higher vimentin expression levels promoted the onset of spheroid invasion. By contrast, the spheroid expansion rates were rather insensitive to changes in matrix confinement except in dense GelMA gels, which completely blocked invasion. The insensitivity of the expansion rate to pore size could potentially be due to cell-mediated matrix remodeling, which may facilitate cell invasion by providing empty space or guiding collagen bundles. The spheroid expansion rate did depend on the MMP1 expression level, suggesting that MMP-mediated matrix degradation aids cell migration and/or cell survival into the matrix. It will be interesting in future studies to analyse the roles of different matrix remodelling mechanisms, including matrix degradation, matrix deposition, matrix crosslinking, and active remodeling by actomyosin-based traction forces. Although matrix stiffness is often described as a factor that influences cell invasion^40, 46^, the invasion measurements in our assays did not demonstrate any clear dependence on hydrogel stiffness. It may be that the initial stiffness of the hydrogels, which we measured by rheology, does not impact the onset of invasion, and that matrix stiffness is altered through matrix remodelling in later stages. Spheroids and invading single cells are known to generate traction forces onto matrices that induce stiffening, while degradation can soften the ECM^47^. Viscoelasticity of matrices is also known to affect collective strand formation and EMT processes^48^. In our hydrogel characterization, measurements of the viscoelasticity of the matrices showed *G^′^ > G^′′^*, indicating that the hydrogels were solid-like. Previous studies of spheroid invasion for MCF-10A and MDA-MB-231 breast cancer cells showed that spheroids can undergo unjamming transitions that depend on cell motility and matrix density^7, 18^. Jammed spheroids do not invade and are considered to be in a solid-like state. Spheroids can unjam to a fluid-like state characterized by collectively invading cell strands or to a gas-like state characterized by dissociation of cells that then migrate individually^17^. In addition to these clearly distinct phases, spheroids can also exist in transitional or in-between phases^18, 49^. To test whether this unjamming framework also applies to MV3 and A549 spheroids, we classified the spheroids as solid-like, fluid-like or gas-like based on the morphological appearance of the spheroids at the end point of the experiments (*t* = 24 hrs for MV3 spheroids, *t* = 72 hrs for A549 spheroids). Since fluid-like spheroid states are characterized by multicellular strands protruding into the matrix, we developed an image analysis method to detect multicellular protrusions and measure their length (Fig. S16). We classified spheroids as fluid-like when the average protrusion length was above 25 µm. We chose this cut-off value since spheroids in their initial state at *t* = 0 occasionally showed apparent protrusions up to this length. Since gas-like spheroid states are characterized by dissociated individually migrating cells, we furthermore measured the number of dissociated cells for all the spheroids. We classified spheroids as gas-like when the individual cell count was above 10. Spheroids with protrusion lengths below 25 µm and no disseminated cells were classified as solid-like. We found that MV3 spheroids sometimes showed co-existence between a liquid-like phase (with long multicellular protrusions) and a gas-like phase (with many dissociated cells), especially in collagen (8 mg/mL) and GelMA (30 mg/mL) (Fig. S17) in control condition. However, when MV3 spheroids were treated with MMP-inhibitor in collagen (8 mg/mL) and GelMA (30 mg/mL), spheroids only existed in a liquid-like phase (cell count below 10) (Fig. S18). Similar analysis was performed for A549 spheroids in control and treated (TGF-*β* and MMP-inhibitor) conditions to determine the different phase transitions in collagen and GelMA hydrogels (Fig. S19).

We then construct a phase diagram of the 3D spheroid invasion states in terms of the degree of matrix confinement (pore size) and the MMP1 expression level of the cells (Fig. 7). Spheroids in matrices with high confinement (small pores) and low cellular MMP1 levels showed solid-like behaviour (blue squares). With increasing MMP1 expression levels, spheroids in collagen (2.4 mg/mL and 8 mg/mL) and GelMA (30 mg/mL) unjammed, initially to a liquid-like state (red squares), and eventually to a gas-like state (yellow squares). Spheroids in GelMA (50 mg/mL) remained solid-like irrespective of the MMP1 expression level. The unjamming phase diagram is qualitatively very similar to earlier unjamming phase diagrams reported for breast cancer cells^7, 18^. However, instead of the ‘cell motility‘ parameter used in earlier studies, we find that the MMP1 expression level is an important determinant of spheroid invasion. The MMP1 expression level depends on cell type (being higher in the mesenchymal-like MV3 cells than in the epithelial-like A549 cells) and it can be tuned in both cell types through TGF-*β* and MMP-inhibitor treatments, which change the levels of MMP1 and other EMT-markers (vimentin and E-cadherin). Because unjamming transitions and EMT both describe the behaviour of immotile multicellular structures that gain motility, the question has been raised in what way these two processes are related^50^. In epithelial monolayers, it was shown that unjamming transitions can operate without EMT^51^. Our findings suggest that in 3D tumoroid invasion, EMT is coupled to unjamming because the transition to a more mesenchymal state enables cells to proteolytically degrade the matrix and thus reduce confinement. Compared to previous experimental studies of spheroid unjamming, which used only collagen gels, we achieved stronger confinement by additionally using GelMA gels with nanometric pores. This allowed us to identify a new regime in the phase diagram where cells that are in principle highly motile (with high MMP1 and vimentin levels) undergo a jamming transition and turn solid-like again. Moreover, MV3 cells under control and TGF-*β* -treated conditions in 30 mg/mL GelMA often showed amoeboid-like behaviour, characterized by cells migrating with a round morphology and blebbing (see Fig. 7, orange square). This behavior suggests the presence of a fourth (or transitional) spheroid phase, in which high matrix confinement pushes highly motile cells towards an amoeboid migration mode before jamming. This behaviour is consistent with described mesenchymal-to-amoeboid transitions of (cancer) cells in response to highly confining microchannels^52, 53^ and micropatterns^14^. It will be interesting to further explore this tentative amoeboid-like migration phase for high confinement and high cell motility in more detail with high-resolution microscopy and carefully tuned GelMA concentrations just below and above 30 mg/mL.

**Figure 7.**
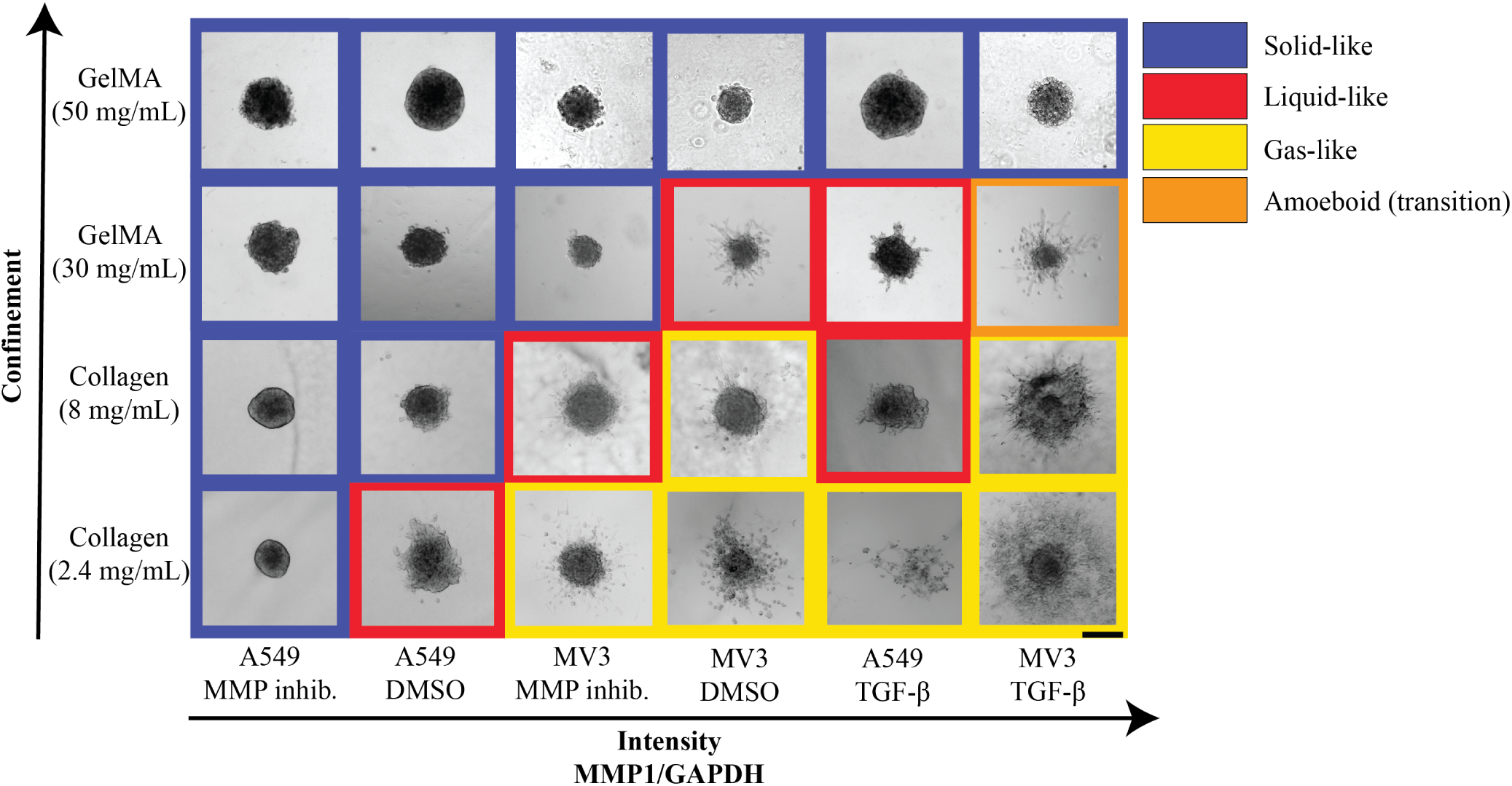
Phase diagram describing (un)jamming transitions of MV3 and A549 cell spheroids in terms of the degree of matrix confinement (which depends on matrix type and concentration) and MMP1 expression level (which depends on cell type and treatment condition). Images were ranked along the x-axis by the MMP1 expression level as quantified by Western blot analysis. Spheroids were classified as gas-like (yellow squares) when more than 10 cells dissociated from the spheroid during the experimental time window, liquid-like (red squares) when the spheroids developed protrusions longer than 25 µm, and solid-like (blue squares) otherwise. Amoeboid behavior (transitional phase) is shown in orange square for MV3 spheroids in GelMA (30 mg/mL). Images are bright-field images taken at *t* = 72 hrs (A549) and *t* = 24 hrs (MV3). Scale bar: 200 µm

Our data are suggestive that TGF-*β* stimulation may drive spheroid unjamming by upregulating the expression level of MMP1. However, we acknowledge that the TGF-*β* pathway has pleiotropic effects and controls many other cellular characteristics^54^. For instance, TGF-*β* has been reported to affect cell-matrix adhesions by downregulating integrins^55^ and to induce cellular traction forces during cell invasion^56, 57^. Confocal reflection microscopy images of MV3 spheroids in collagen (2.4 mg/mL) at the endpoint of invasion experiments (*t* = 24 hrs) indeed indicated that TGF-*β* treatment not only enhanced matrix degradation, but also activated cellular traction forces as evidenced by densified and aligned collagen fibers (Fig. S20). We note that the collagen images also revealed that the MV3 spheroids exerted larger traction forces on the matrix than A549 spheroids, as indicated by radially oriented collagen fibers around the MV3 spheroids as opposed to more uniform collagen networks around A549 spheroids^58^. These differences in the mechanical cell-matrix interplay could also have impacted the invasion, as tensile forces on collagen are known to influence the onset of invasion^59^. To our surprise, MMP inhibition in the epithelial-like A549 cells resulted in reduced E-cadherin levels compared to control conditions. MMPs, including MMP1 and MMP2, have been shown to induce EMT by regulating E-cadherin levels^60–62^. Several MMPs are associated with E-cadherin cleavage and the soluble E-cadherin extracellular domain that results from cleavage was found to promote cancer invasion by in turn increasing MMP production^61, 63^. MMP-mediated E-cadherin cleavage is also found downstream of TGF-*β* activation, providing a pathway for reduced E-cadherin levels in TGF-*β* -treated A549. However, the downregulation of E-cadherin levels in response to MMP inhibition was unexpected, and to the best of our knowledge an effect that has not previously been reported. We propose this effect could be an interesting direction for future research to further elucidate the mechanisms behind MMP inhibitors, which so far failed in the clinic for cancer treatments^61^.

In this work, we did not distinguish between invasion by cell migration versus cell proliferation. Proliferation could play a role in setting the onset time for invasion, as proliferation can increase spheroid packing densities and increase the active forces of cells that regulate solid-to-liquid transitions^64^. Furthermore, the initiation of invasion has been linked to solid tumor growth^65^. Tumor proliferation and invasion are interconnected processes and changes in proliferation could have impacted the normalized spheroid radius over time^66^. To improve our understanding of the contribution of proliferation in invasion, follow-up studies could measure proliferation rates or proliferation could be blocked by adding chemical agents or cell synchronization^67^, although these agents might cause undesired effects on the cells, e.g. by causing replication stress or increased cell death^68^.

While jamming-unjamming transitions have become a well-established framework to explain cancer invasion, the underlying biological mechanisms are poorly understood. Recent studies have revealed possible mechanisms that impact plasticity pathways such as the ERK1/2 pathway, which triggers cell motility and results in collective invasive strands^16^, and a mechanosensitive pathway, in which cellular confinement directly causes deformations of the cell nuclei accompanied by chromatin remodelling^69^. Here we showed evidence that EMT is also a mechanism involved in 3D unjamming transitions by regulating cell-matrix interplay. This is in line with earlier findings that reported EMT-independent unjamming transitions in 2D monolayers^51^. Our findings demonstrate that the vimentin expression level impacts the onset time of spheroid unjamming, while the MMP1 expression level impacts the subsequent spheroid expansion rate and associated unjamming transitions. Understanding the biological mechanisms behind unjamming transitions is beneficial for cancer patients, as unjamming is considered a predictor for distant metastasis^70^.

## Methods

### Cell culture

MV3 human melanoma cells that were three times xenografted in nude mice and selected for highly-metastatic behaviour (a kind gift from Peter Friedl,^71^), were cultured in DMEM-F12 (Dulbecco’s Modified Eagle Medium, Thermo Scientific) supplemented with 10% Fetal Bovine Serum (FBS, Thermo Fisher) and 1% penicillin/streptomycin (Thermo Fisher). A549 human lung adenocarcinoma cells (CCL_185, ATCC) were cultured in DMEM (Gibco, #41965039), and supplemented with 10% FBS (Gibco) and 1% Antibiotic-Antimycotic solution (Gibco). All cells were incubated at 37°C and 5% CO_2_, passaged at 80-90% confluency, and sub-cultured 2-3 times per week.

### Spheroid formation

Spheroids were grown in a commercially available Corning*^TM^*Elplasia*^TM^* 96-well plate for high-throughput spheroid production. These well plates are round-bottom with an Ultra-Low Attachment (ULA) surface that prevents cell-surface attachment and promotes cell-cell adhesion. We used an initial seeding density of 40 x 10^3^ cells (500 cells per micro-well) for each well to produce 79 spheroids. Spheroids were ready to use after 4 days of culture in the wells and had an average diameter of approximately 200 *±*30 µm for both cell lines (Fig. S21). We restricted the spheroid diameter to less than 250 µm to avoid a necrotic core.

### Hydrogel preparation

Bovine hide collagen type I (purity *≥* 99.9%, Advanced Biomatrix) stock solutions (3 mg/ml and 10 mg/ml in 0.01 N HCL) were used to prepare 2.4 mg/ml and 8 mg/ml collagen gels, respectively. The collagen was made isotonic by adding 12.5 v/v% of 10x Phosphate Buffered Saline (PBS, Thermo Fisher) to collagen. In addition, 0.1 M sodium hydroxide was added to bring the pH to 7.4. All solutions were kept on ice. Pre-cooled Milli-Q (MQ) was added to bring the final collagen concentration to either 2.4 or 8 mg/mL. Final collagen dilutions were vortexed for 30 seconds and polymerized at 37°C in an *µ*-slide 8-well (Ibidi) for at least 45 minutes. Gelatin methacryloyl (GelMA) was purchased from Sigma Aldrich (300g bloom, 60% degree substitution). GelMA retains the thermo-reversibility of gelatin^72^, but the methacrylic anhydride groups can undergo covalent cross-linking under UV light (365 nm) in the presence of a photoinitiator. We used 3 and 5wt% GelMA with a 1:16 mass ratio of photoiniator (Lithium phenyl-2,4,6-trimethylbenzoylphosphinate, LAP; Sigma Aldrich). LAP and GelMA were dissolved together in Dulbecco’s Phosphate Buffered Saline (DPBS; Gibco) at 37°C in a water bath for about 2 hrs. The hydrogel was crosslinked using a UV-lamp (Spectroline, Serial no. 1832066) at a wavelength of 365 nm for 45 s.

### 3D invasion assays

Spheroids were embedded in collagen gels by a sandwich protocol, adapted from^73^. First, collagen gel layers of 80 µL were polymerized in *µ*-Slide 8 well chambers (ibidi) for 45 minutes at 37°C. Next, spheroids were transferred from the culture plate onto the collagen layers by carefully pipetting them in culture medium. Spheroids were incubated on the collagen gel for 30 minutes to ensure attachment. Next, culture medium was pipetted out of the chambers and collagen layers of 100 µL were added on top of the spheroids and polymerized for 45 minutes at 3°C. After polymerization, 200 µL of cell culture medium was added to the spheroid-collagen gels, which were then incubated at 37°C and 5% CO_2_. For spheroids embedded in GelMA, we pipetted 125 µL of GelMA/spheroid suspension in a well of the *µ*-Slide 8-well chambers (ibidi). We then exposed the *µ*-Slide to 45 s of UV-light (365 nm) using a lamp, followed by addition of 200 µL culture medium.

Recombinant TGF-*β* (stock concentration 5 µg/mL was diluted in culture medium to a final concentration of 10 ng/mL^74, 75^. Batimastat (BB-94) broad spectrum MMP-inhibitor (stock concentration 1 mM) from Abcam was diluted to 30 µ in culture medium^76^. The supplemented media were added to the respective ibidi wells with spheroids embedded in hydrogels before incubation and imaging. Any cancer spheroids that made contact with the glass substrate or the side walls of the *µ*-Slide (ibidi) were excluded from analysis. Spheroid invasion was monitored using a Colibri Axio Observer 7 inverted microscope under bright-field settings with a 5x/NA 0.16 air objective for time-lapse imaging (multiple positions) at a time interval of 1 hr. MV3 spheroid invasion imaging was restricted to 24 hrs, while A549 spheroid invasion was imaged until 72 hrs. All experiments were conducted at 37°C, 90% humidity and 5% CO_2_ using a stage top incubator (ibidi).

### Rheology

Shear rheology of hydrogels was performed on a Kinexus pro+ rheometer (Malvern, UK), using a 20 mm stainless steel parallel plate and 0.5 mm gap. The collagen samples were pipetted on the bottom plate, which was held at 4°C using a Peltier system, the top plate was immediately lowered, and the temperature was set at 37°C to induce collagen polymerization. The GelMA samples were pipetted on the bottom plate at 37°C and were crosslinked using a UV lamp (Spectroline, Serial no. 1832066) for 45 seconds at 365 nm wavelength at a fixed distance of 2 cm from the sample. Immediately after crosslinking, the top plate was lowered. Mineral oil was added around the sample edge to prevent solvent evaporation. During and after polymerization, small amplitude oscillatory shear was applied to the hydrogels with a constant strain amplitude (0.5%) and frequency (1Hz) to measure the storage modulus (*G*’) and loss modulus (*G*”).

### Image and data analysis for 3D invasion assays

We developed a custom-made MATLAB script to detect the spheroid boundary and disseminated single cells in bright-field time-lapse image series of invading spheroids. We first adjusted the image brightness and contrast using the MATLAB command ’imsharpen’. We then created a binary gradient mask by adjusting the image segmentation threshold to detect discontinuities in brightness using the derivative of a Gaussian filter. This threshold value was optimized for each set of bright-field images. The binary gradient mask outlined the identified object, which was then post-processed to smoothen and dilate using the MATLAB command ‘imdilate‘. This dilated gradient mask finally underwent the ’fill hole’ process using the MATLAB command ‘imfill‘. The spheroid boundary (outlined in red in Fig. S9) was detected by MATLAB command ‘bwboundaries‘ that identifies exterior boundaries. From the boundary, we calculated the spheroid area and effective circular radius. To account for variations in initial spheroid size, we normalized the effective circular radius by its initial value at *t* = 0. Single dissociated cells were identified with the same threshold parameters (highlighted in green, see Fig. S9). In case individual cells were already present at *t* = 0, we subtracted these from the cell count at the end of the assay (24 hrs for MV3, 72 hrs for A549). Note that the cell counts likely underestimate the actual cell number since the limited contrast of the images made it difficult to distinguish whether cells close to the spheroid boundary had dissociated.

To identify multicellular protrusions of MV3 and A549 spheroids and determine their lengths, we developed another MATLAB script. First we converted 2D bright-field images from the Cartesian coordinate system (*x*,*y*) to a polar coordinate system (*θ*, *r*). Next we detect protrusions as peaks relative to the average spheroid radius (highlighted in cyan, see Fig. 16C, D). The protrusion length analysis was performed on images obtained at the end time point of each experiment (*t* = 24 hrs for MV3, *t* = 72 hrs for A549). The average protrusion length was used as a criterion to distinguish between a solid-like phase (length below 25 µm) versus a liquid-like state (length above 25 µm).

### Western Blot analysis

Bovine type I collagen (Procol, Advanced Biomatrix, 3 mg/ml) was diluted 1:100 with MilliQ water and pipetted into six-well plates (Thermo Fisher) to cover the surface and left to incubate for 2 hours at room temperature. The collagen coatings were washed twice with PBS. Next, MV3 cells (300,000/well) and A549 cells (400,000/well) were seeded in the coated plates in culture medium under control conditions 2 µL/mL DMSO) or with recombinant TGF-*β* at 10 ng/mL (stock concentration 5 ng/*µ*L) or Batimastat at 30 *µ*M (stock concentration 1 mM, BB-94/Abcam). After 48 hours (or 1 hour for SMAD activity tests), cells were washed with PBS, lysed with cold radioimmunoprecipitation buffer (RIPA, 100 µL/well, Thermo Fisher) and transferred to Eppendorf tubes. Lysed samples were agitated at 4°C for 30 minutes and stored at -20°C. Laemmli buffer (2x, Bio-rad) and 4% *β* -mercaptoethanol (Sigma Aldrich) were added to the lysed samples, which were incubated at 95°C for 5 minutes. Sodium dodecyl sulfate-polyacrylamide gel electrophoresis (SDS-PAGE) was performed with Mini-PROTEAN TGX gels (Bio-rad) using 100V for 1.5 hours. Western Blotting was executed with a Trans-Blot Turbo Transfer System (Bio-rad) and Trans-Blot Turbo Mini 0.2 µm PVDF Transfer Packs (Bio-rad). The membranes were blocked in 5% Bovine Serum Albumin (BSA, Thermo Fisher) in phosphate buffered saline (PBS, company) overnight. Membranes were stained with primary antibodies: rabbit anti-SMAD2 (1:1000, #5339, Cell signaling), rabbit anti-pSMAD2 (1:1000, kind gift from Peter ten Dijke^77^), rabbit anti-MMP1 (1:1000, #54376, Bioke), mouse anti-MMP2 (1:1000, #436000, Thermo Fisher), rabbit anti-E-cadherin (1:1000, #ab40772, Abcam), mouse anti-vimentin (1:2000, #ab8978, Abcam) and rabbit anti-GAPDH (#CST2118S, Bioke) in 5% BSA overnight on a shaker at 4°C. Membranes were washed thrice with 0.1% Tween (Sigma Aldrich) in PBS (PBS-T) on a shaker, and incubated for 3-5 hours with secondary antibodies: rabbit anti-mouse HRP (#ab97051, Abcam) and goat anti-rabbit HRP (#ab6728, Abcam), 1:5000 in PBS-T. Afterwards, membranes were washed thrice with PBS-T and imaged with an enhanced luminol-based chemiluminescent substrate kit (Thermo Fisher) on a gel imager (Bio-rad).

### Pore size and void fraction analysis of collagen and GelMA hydrogels

Collagen and GelMA hydrogels were prepared in *µ*-Slide 8 well chambers and were imaged by confocal reflectance on a Stellaris 8 confocal microscope (Leica), equipped with a white light laser and a hybrid detector (HyDS), using a 488 nm laser line and a 63x magnification objective (1.3 NA, glycerol immersion) at room temperature. We recorded Z-stacks starting at 100 µm above the glass surface over a total depth of 30 µm with a 2 µm step size. For the collagen networks, we used a custom-made python script^78, 79^, which implements the bubble method to measure the sizes of the pores in between collagen fibers (see Fig. S2.

Confocal images were denoised using the total variation minimization method^80^ on a slice-by-slice basis. A local threshold was then applied to obtain a binary confocal stack. The Euclidean distance map was determined, representing the distance to the nearest fiber at each point in the image, and a Gaussian filter was applied with a standard deviation of 5 pixels to the Euclidean distance map. Finally, the local maxima of the Euclidean distance map were determined, which represent the furthest distance from a fiber in the image. We selected these distances as half of the pore size. The average pore size was calculated of three replicates for each concentration (n=3). The GelMA hydrogels with their nanometric pores were too dense to be able to directly measure the pore size distribution. Instead, we performed a void fraction analysis on maximum intensity projections of confocal reflectance microscopy Z-stacks of GelMa and collagen hydrogels made in Fiji^81^. The images were adjusted for brightness and contrast by maximum intensity projection. The adjusted images were then converted to a binary image using the ‘Make Binary‘ function in Fiji with white (1) pixels corresponding to hydrogel fibers and black (0) pixels corresponding to the voids. We determined the void fraction as the ratio of black pixels to white pixels.

### GelMA hydrogel permeability analysis

Permeability analysis was performed on GelMA hydrogels (30 and 50 mg/mL) to estimate their pore size. The permeability, *K* of a hydrogel can be obtained from Darcy’s Law by estimating the flow velocity of tracer particles through the material. To do this, we used a microfluidic platform equipped with three channels, where the middle channel was filled with GelMA hydrogel and crosslinked by a UV-light source (365 nm)^75^. The top and bottom channels were operated using a pressure pump device to create a pressure gradient that drives the flow through the hydrogel channel at room temperature. The microfluidic chip was mounted on Zeiss Axio Observer 7 equipped with an ORCA Flash 4.0 V2 (Hamamatsu) digital camera with a resolution of 2048 *×* 2048 pixels. We used Rhodamine B dye (Merck Sigma) (1% v/v solution in 1X Dulbecco’s Phoshate Buffer Solution from Sigma Aldrich) as a tracer particle. After applying a pressure gradient ΔP = 20 mbar, the displacement of the dye front was tracked by time-lapse imaging with a time interval of 30 seconds. Imaging was performed using 543 nm LED laser (excitation/emission: 543 nm/568 nm) at 30% intensity and 1.58 s exposure time, and a 5x (NA 0.16/Air) objective. The images were then and converted into 8-bit color images using ImageJ^81^. A line was drawn in the direction of the flow to obtain a histogram of the fluorescence intensity as a function of distance. The fluorescence intensity is tracked by identifying the distance (pixel value *×* pixel length (0.62 µm) value at which the mean intensity of fluorescence is zero. This position was recorded for the next 10 consecutive images at 30 second interval. The difference between two consecutive images is divided by the time interval between images to obtain flow velocity. Averaging for each consecutive images thus gives the average flow velocity. From the measured velocity, we computed the hydraulic permeability of the GelMA hydrogels using the Darcy equation:

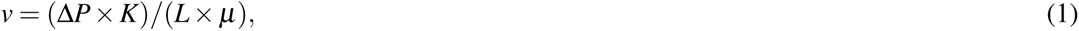

where *µ* is the fluid velocity across the hydrogel (m/s), ΔP is the pressure gradient across the hydrogel (Pa), *L* is the length of the hydrogel channel in the direction of the flow (m), *K* is the hydraulic permeability (m^2^), and *µ* is the fluid dynamic viscosity (10*^−^*^3^ Pa.s). We recovered an average permeability of 2.36 *×* 10*^−^*^16^ m^2^ for 30 mg/mL GelMA and 1.29 *×* 10*^−^*^16^ m^2^ for 50 mg/mL GelMA. The corresponding pore sizes, calculated as the square root of the permeability, were 1.54 nm for 30 mg/mL GelMA and 1.14 nm for 50 mg/mL GelMA (for a detailed summary see Table S1).

## Acknowledgements

A.V.D.N. and G.H.K. gratefully acknowledge funding from the OCENW.GROOT.20t9.O22 project *The Active Matter Physics of Collective Metastasis* financed by the Dutch Research Council (NWO). Z.R. and P.E.B gratefully acknowledge funding from the European Research Council (ERC) under the European Union’s Horizon 2020 research and innovation program (grant agreement no. 819424). P.T.D. and P.E.B. are supported by ZonMW grant (09120012010061). A.B. gratefully acknowledges funding from MSCA Postdoctoral Fellowships 2022 Project ID: 101111247. I.M. gratefully acknowledges funding from the Convergence programme Syn-Cells for Health(care) under the theme of Health and Technology.

## Author contributions statement

Conceptualization, A.V.D.N., Z.R., P.E.B. and G.H.K.; Investigation, A.V.D.N. and Z.R.; Formal analysis, A.V.D.N., Z.R., A.B. and I.M.; Software, Z.R., A.B. and I.M.; Visualization: A.V.D.N. and Z.R.; Writing – Original Draft, A.V.D.N. and Z.R.; Writing – Review & Editing A.B., I.M, P.T.D., P.E.B. and G.H.K.; Funding acquisition, P.E.B. and G.H.K.; Supervision, P.E.B. and G.H.K.

