## Supplemental file for "EMT-dependent cell-matrix interactions are linked to unjamming transitions in cancer spheroid invasion"

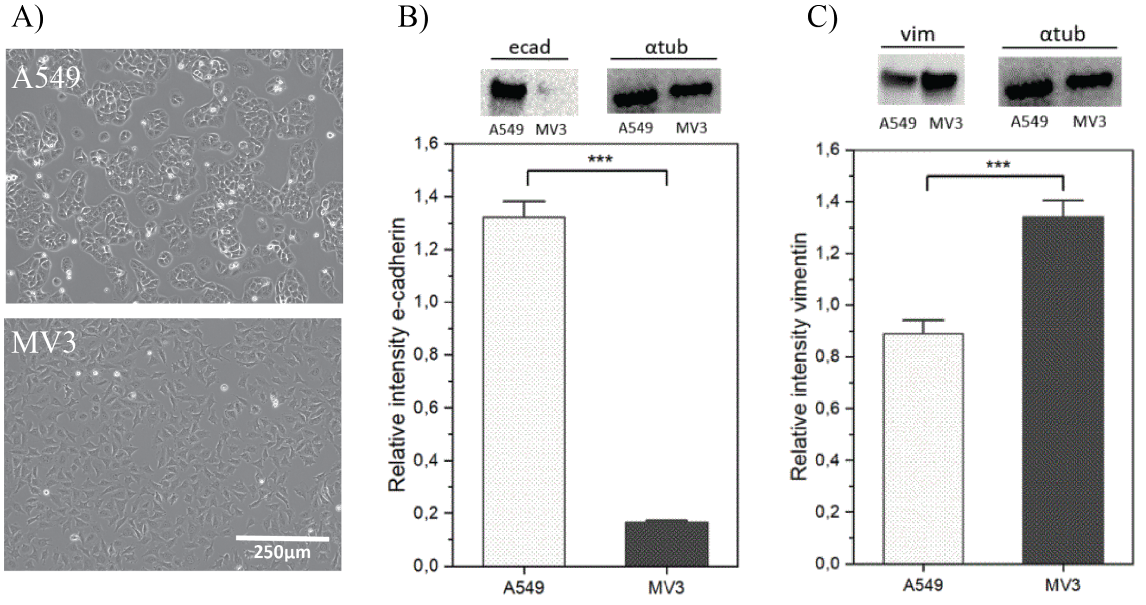


Figure 1: **Cell morphology and Western Blot analysis for A549 and MV3 cancer cells.** A) Bright-field images of the cells cultured on collagen-coated 6-well dishes. A549 cells (top panel) show clustering, which is a characteristic of cells expressing high levels of E-cadherin (epithelial marker). MV3 cells (bottom panel) show individual spindle-shaped cells with mesenchymal-like features. Scale bar: 250 $\mu$m. B) Western Blot analysis of A549 and MV3 cells for E-cadherin (ecad) expression levels, normalized to $\alpha$-tubulin expression, showing that A549 cells are more epithelial-like than MV3 cells. (C) Corresponding Western Blot analysis for vimentin expression levels, showing that MV3 cells express significantly more vimentin than A549 cells.

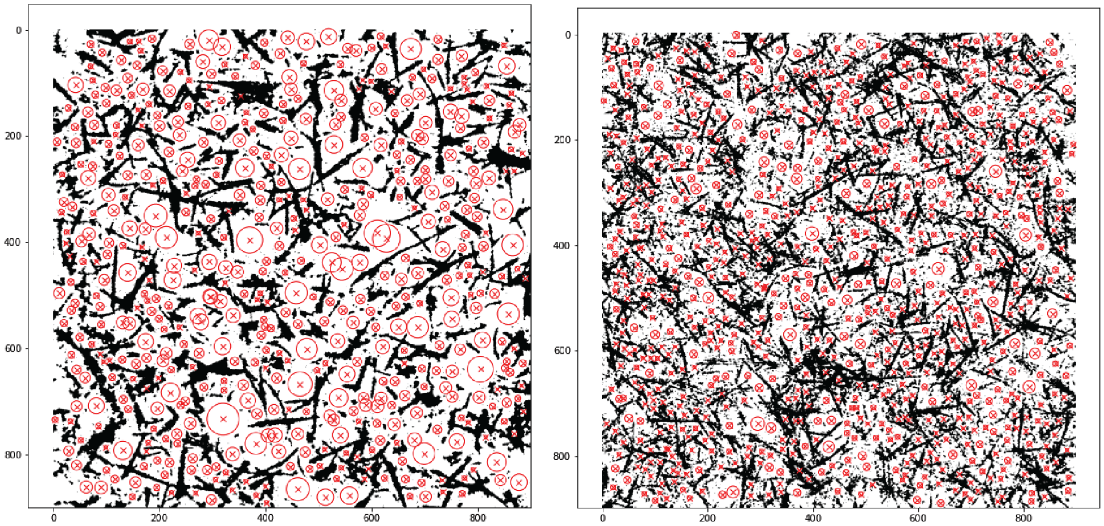


Collagen (2.4 mg/mL) Collagen (8 mg/mL)

Figure 2: Binarized confocal microscopy reflection images of collagen networks with concentrations of 2.4 mg/mL (left) and 8 mg/mL (right) with fitted bubbles (red circles), to determine the pore size distribution. The x- and y-axes indicate the length scale in pixels.

| Repeat | Average velocity | Hydraulic permeability ($m^2$) | Pore size (nm) |
|---|---|---|---|
| GelMA (30 mg/mL) | | | |
| 1. | 0.54 | $2.43 \times 10^{-16}$ | 15.58 |
| 2. | 0.51 | $2.29 \times 10^{-16}$ | 15.13 |
| 3. | 0.53 | $2.38 \times 10^{-16}$ | 15.42 |
| GelMA (50 mg/mL) | | | |
| 1. | 0.28 | $1.26 \times 10^{-16}$ | 11.22 |
| 2. | 0.32 | $1.44 \times 10^{-16}$ | 12.00 |
| 3. | 0.26 | $1.17 \times 10^{-16}$ | 10.81 |

Table 1: **Average pore size estimated from hydraulic permeability measurements for GelMA hydrogels with concentrations of 30 mg/mL (15.3 $\pm$ 0.18 nm) and 50 mg/mL (11.3 $\pm$ 0.5 nm).** The permeability was determined from the average flow velocity of Rhodamine B dye measured in permeation experiments performed in a microfluidic chip at a pressure gradient of 20 mbar.

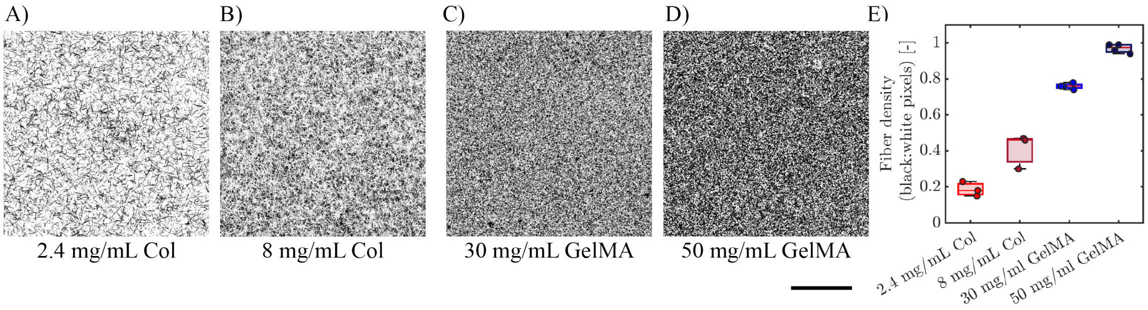


Figure 3: **Hydrogel fiber density measurements based on confocal reflectance images.** Binarized maximum intensity projections of confocal Z-stacks composed of 16 slices at 2 $\mu$m intervals for hydrogels of A) collagen (2.4 mg/mL), B) collagen (8.0 mg/mL), C) GelMA (30 mg/mL) and D) GelMA (50 mg/mL). E) Fiber density (black to white pixel ratio) measurements for hydrogels. Data are averages with standard deviations for 3 repeats. Scale bar: 25 $\mu$m.

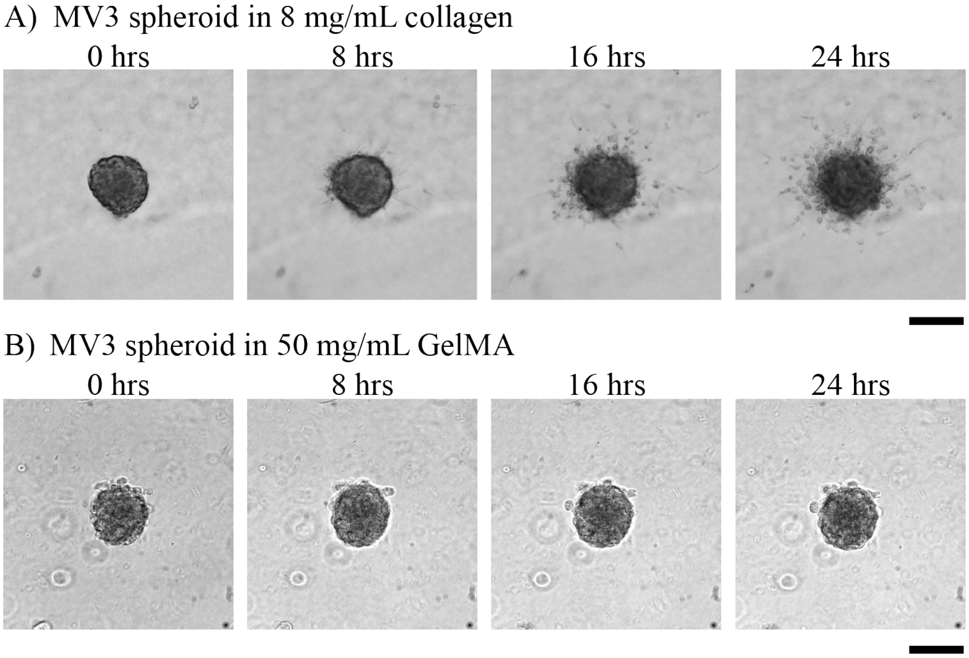


Figure 4: **MV3 spheroid invasion in collagen and GelMA matrices.** Bright-field images of MV3 spheroids at 8 hr time intervals in A) 8 mg/mL collagen and (B) 50 mg/mL GelMA at t=0, 8, 16 and 24 hrs. Scale bars: 200$\mu$m. MV3 spheroids invaded into the collagen network, as seen from the spheroid protrusions and presence of disseminated single cells, but did not invade in the GelMA hydrogel.

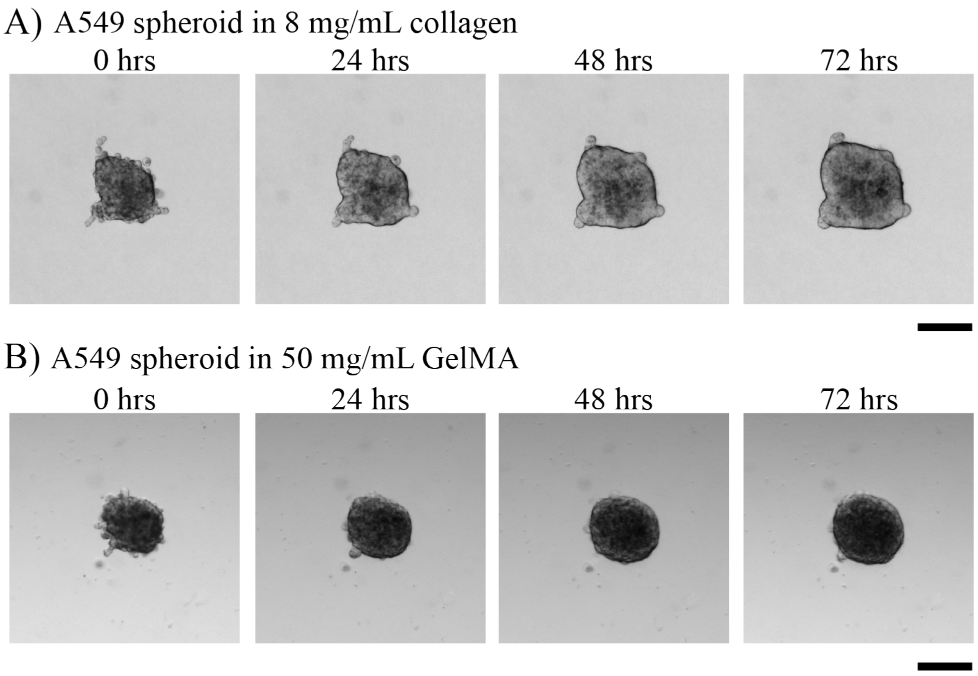


Figure 5: **A549 spheroid invasion in collagen and GelMA matrices.** Bright-field images of A549 spheroids at 24 hr time intervals in A) 8 mg/mL collagen and (B) 50 mg/mL GelMA. Scale bars: 200$\mu$m. The A549 spheroids had an irregular shape with occasional protrusions. Over time, the spheroid boundary remained rather smooth and the spheroids uniformly expanded in size without the formation of protrusions or dissociation of single cells.

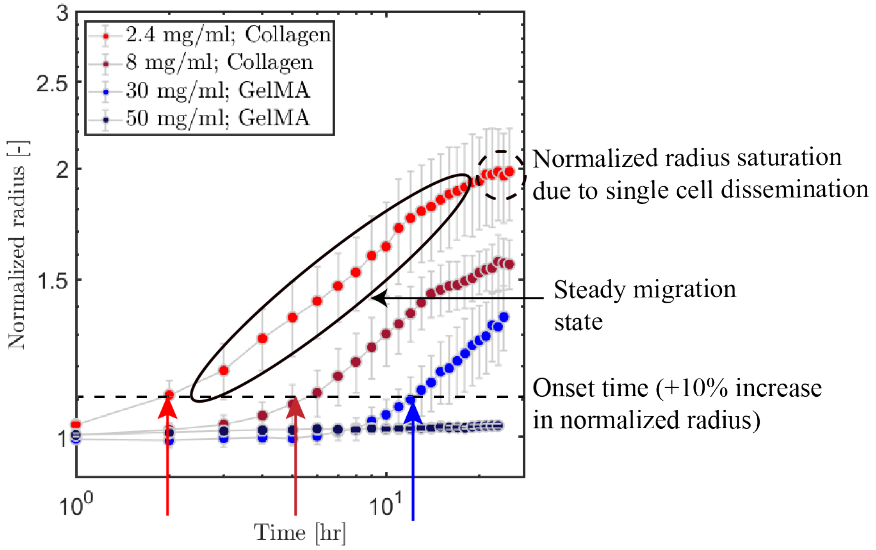


Figure 6: **Definition of invasion onset time and spheroid expansion, illustrated for MV3 spheroids.** The effective circular radius of MV3 spheroids normalized by its initial value is tracked as a function of time for spheroids in collagen and GelMA hydrogels (see legend). The invasion onset time is defined as the time point where the normalized radius reaches a value of 1.1 (indicated by the arrows and the horizontal dashed line). The expansion rate was defined as the power-law slope of the spheroid growth curves after the onset of invasion. Since the growth curves plateaued at long time due to the dissociation of cells from the spheroid (dashed circle), the curves were fitted within the time range enclosed within the solid ellipse. The maximum time for the fit range was taken as the time point where the difference in normalized spheroid radius became less than 0.01 between two successive time points.

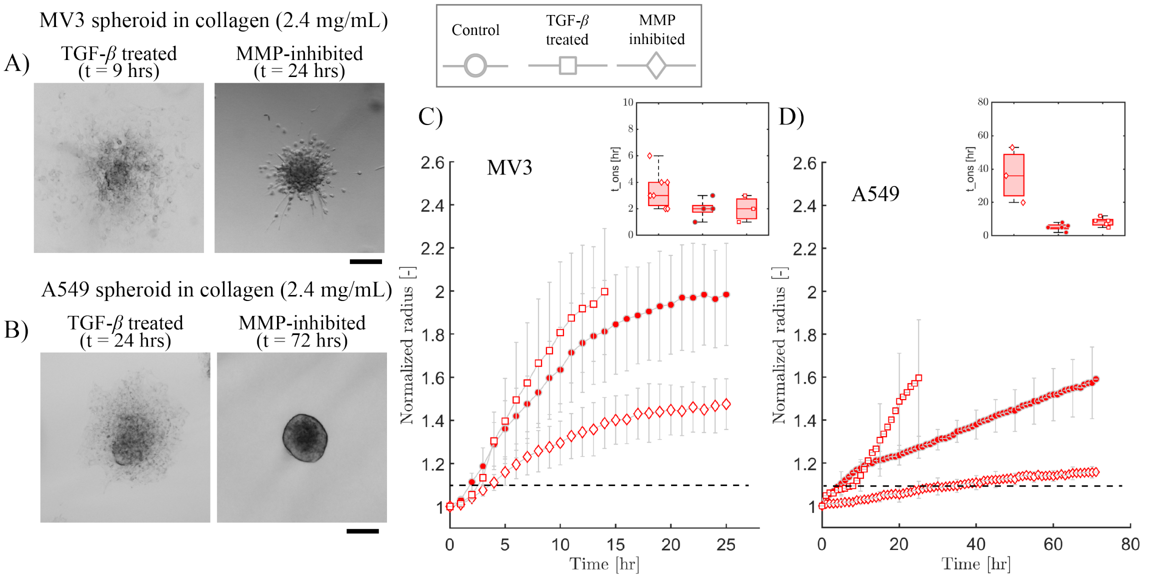


Figure 7: **Effect of TGF-$\beta$ and MMP-inhibitor treatments on spheroid invasion in collagen (2.4 mg/mL).** A) Bright field images of MV3 spheroids treated with TGF-$\beta$ (left, t = 9 hrs) and with MMP-inhibitor (right, t = 24 hrs). B) Bright field images of A549 spheroids treated with TGF-$\beta$ (left, t = 24 hrs) and MMP-inhibitor (right, t = 72 hrs). Scale bars in A,B: 200 $\mu$m. C) Increase in spheroid radius with time for MV3 spheroids under control, TGF-$\beta$, and MMP inhibition conditions (see legend). D) Corresponding data for A549 spheroids. Horizontal dashed lines in C,D denote the threshold value for the normalized radius of 1.1 defined as the onset of invasion. Insets show the invasion onset times in control and treated conditions. Neither treatment affected the onset time of invasion much, except for MMP-inhibited A549 spheroids, where invasion was drastically delayed. Note that upon TGF-$\beta$ treatment, the spheroid cores eventually disintegrated and sedimented to the bottom glass surface. In this condition we therefore only tracked invasion up to t = 14 hrs for MV3 spheroids and up to t = 24 hrs for A549 spheroids.

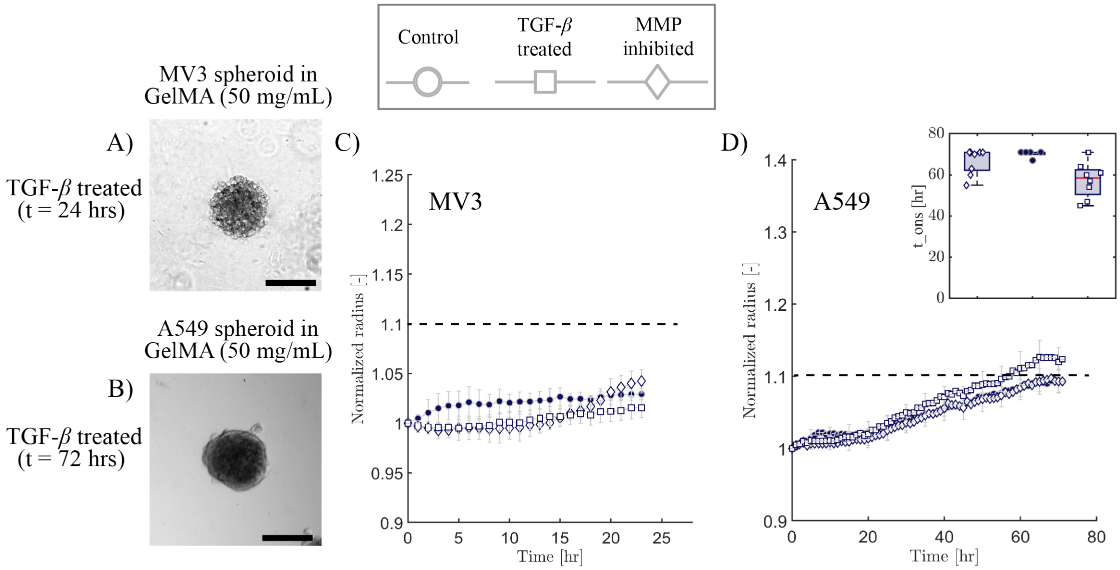


Figure 8: **Effect of TGF-$\beta$ and MMP-inhibitor treatments on spheroid invasion in GelMA (50 mg/mL).** A) Bright field image of a MV3 spheroid treated with TGF-$\beta$ at t = 24 hrs. B) Bright field image of an A549 spheroid treated with TGF-$\beta$ at t=72 hrs. Neither the MV3 nor the A549 spheroids showed protrusions or single cell dissemination, despite the TGF-$\beta$ treatment. Scale bars in A, B: 200 $\mu$m. (C) Increase in spheroid radius with time in control, TGF-$\beta$ treated and MMP-inhibited conditions (see legend on top) for C) MV3 spheroids and D) A549 spheroids. Horizontal dashed lines in C,D denote the threshold value for the normalized radius of 1.1 defined as the onset of invasion. Inset in D: onset time of invasion for A549 spheroids. TGF-$\beta$ treated A549 spheroids had a slightly lower onset time (at t=57 hrs) compared to control and MMP-inhibited conditions (at t=68 hrs). Note that we could not determine the onset of invasion for the MV3 spheroids over the 24 hrs experimental time frame.

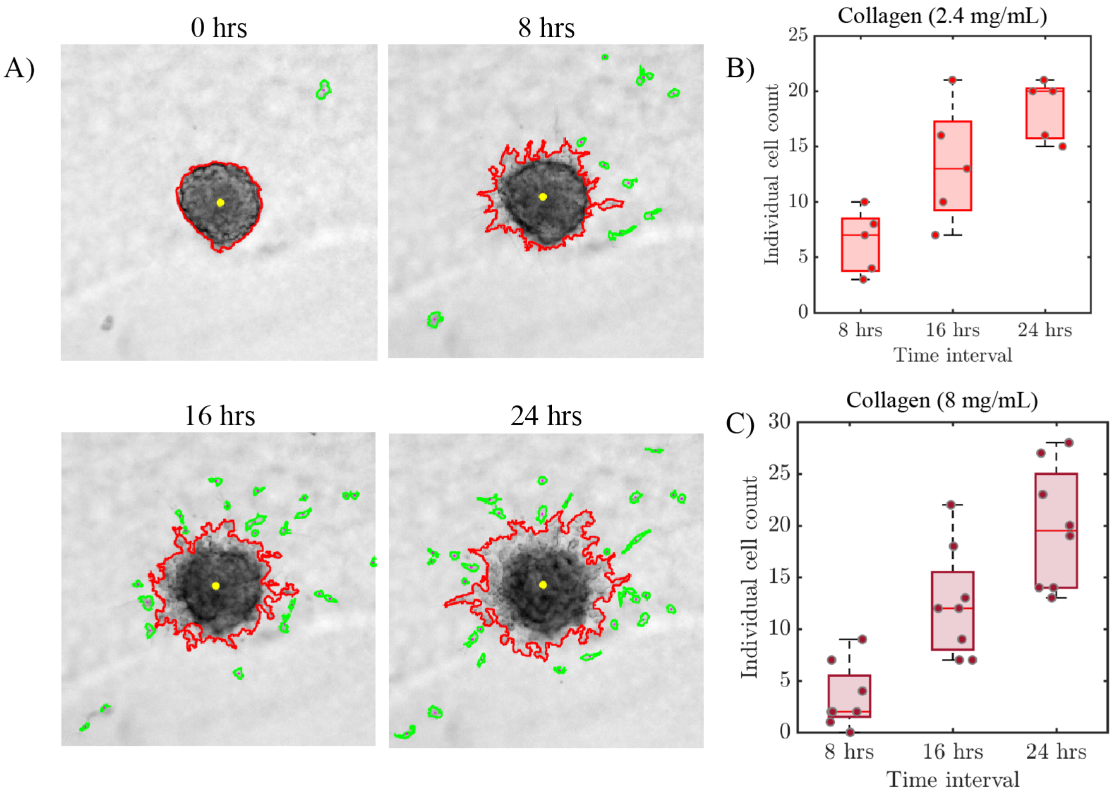


Figure 9: **Quantification of cell dissociation from MV3 spheroids in collagen matrices.** A) Bright-field images of MV3 spheroids in collagen (8 mg/mL) at 8 hr time intervals. The spheroid border is outlined in red while dissociated cells are outlined in green. B) Number of individual cells disseminated in 2.4 mg/mL collagen at time intervals of t=8, 16 and 24 hrs. C) Corresponding data for MV3 spheroids in 8 mg/mL collagen.

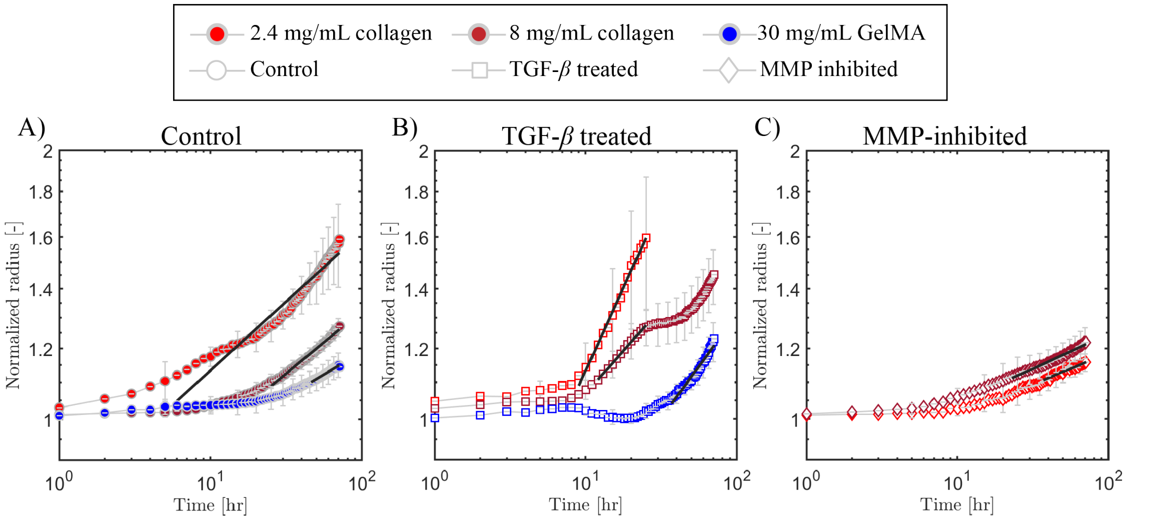


Figure 10: **Effect of TGF-$\beta$ and MMP-inhibitor treatments on expansion rates of A549 spheroids.** Quantification of the increase in spheroid radius with time for A549 spheroids in collagen (2.4 mg/mL and 8 mg/mL) and GelMA (30 mg/mL) under A) control conditions, B) with TGF-$\beta$ treatment, and (C) with MMP-inhibitor treatment. The expansion rates were determined as the power law slope of the curves after the onset of invasion. The solid black lines indicate the fits together with the temporal range.

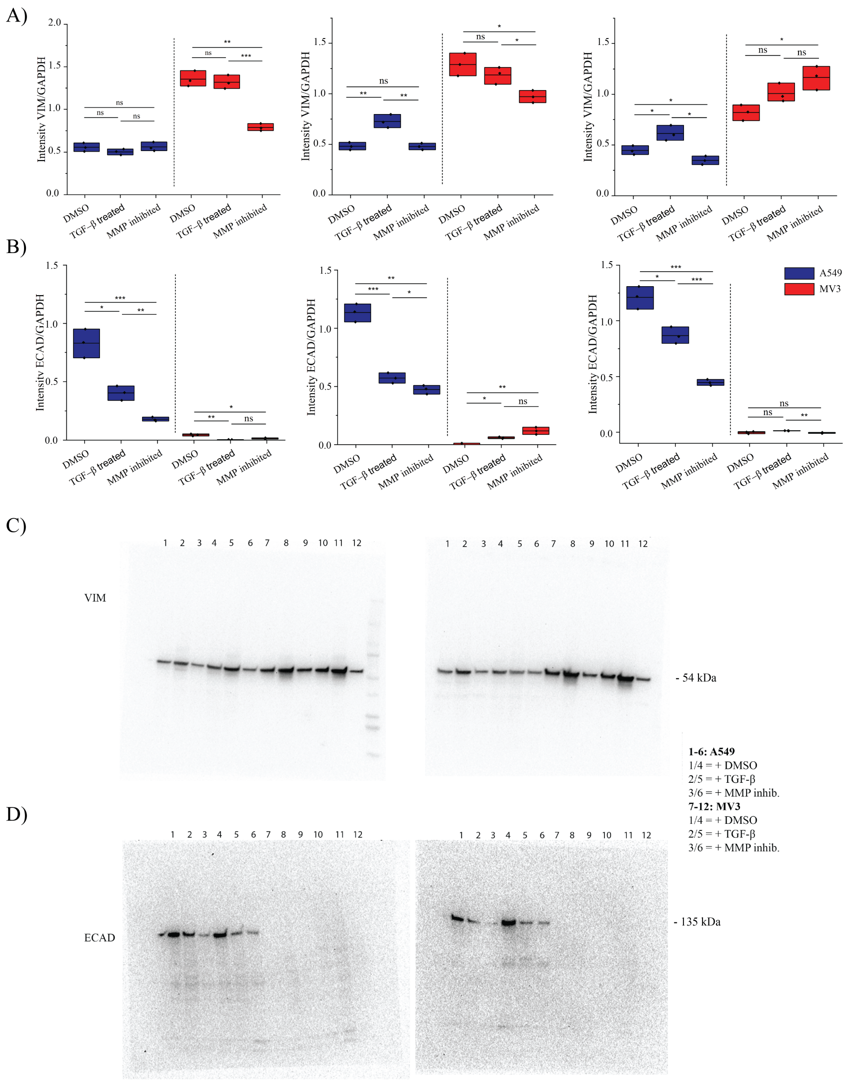


Figure 11: **Western Blot analysis for vimentin and E-cadherin protein expression levels in A549 and MV3 cells cultured in 2D on collagen-coated 6-well plates.** We performed a total of n = 4 biological replicates for each condition (DMSO (control), TGF-$\beta$, and MMP-inhibitor treatments). One replicate of each is shown in the main text. Additional replicates are shown here. A) Three replicates showing vimentin expression levels normalized by GAPDH. B) Three replicates showing E-cadherin expression levels normalized by GAPDH. Two examples of Western Blot images are shown for C) E-cadherin and D) vimentin. The legend on the right shows the content of each lane (indicated by numbers above the gels).

A)

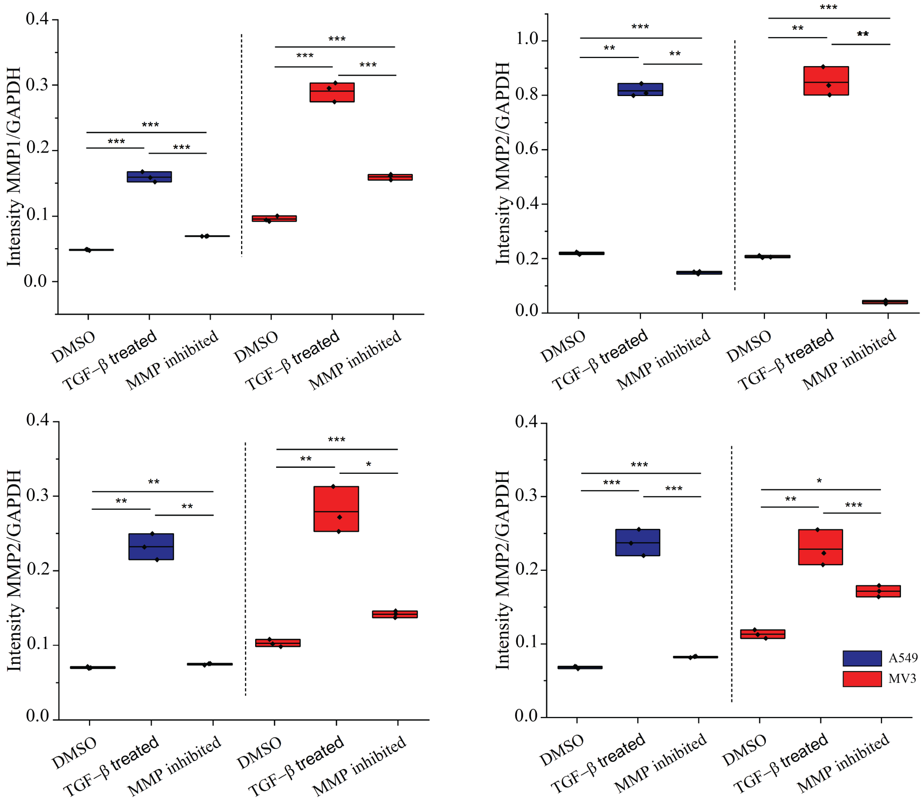


B)

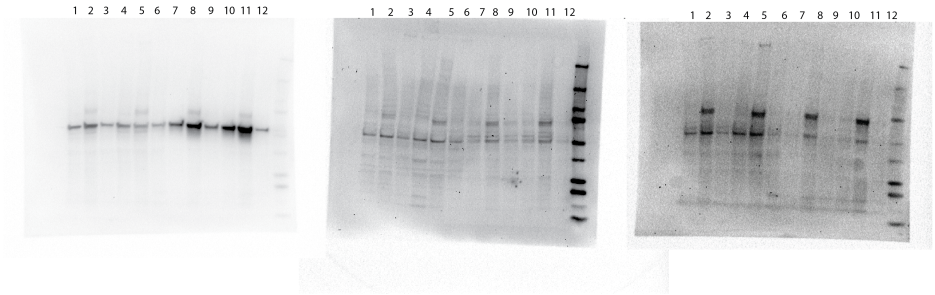


Figure 12: **Western Blot analysis for MMP1 and MMP2 protein expression levels in A549 and MV3 cells cultured in 2D on collagen-coated 6-well plates.** We performed a total of n = 2 biological replicates for the MMP1 quantification and n = 4 for the MMP2 quantification, for different conditions: DMSO (control), TGF-$\beta$, and MMP-inhibitor treatments. One replicate of each is shown in the main text. Additional replicates are shown here. A) One replicate showing MMP1 expression levels normalized by GAPDH and three replicates showing MMP2 expression levels normalized by GAPDH. C) Three Western Blot images are shown for MMP1 (left) and MMP2 (middle and right). The legend on the right shows the content of each lane (indicated by numbers above the gels).

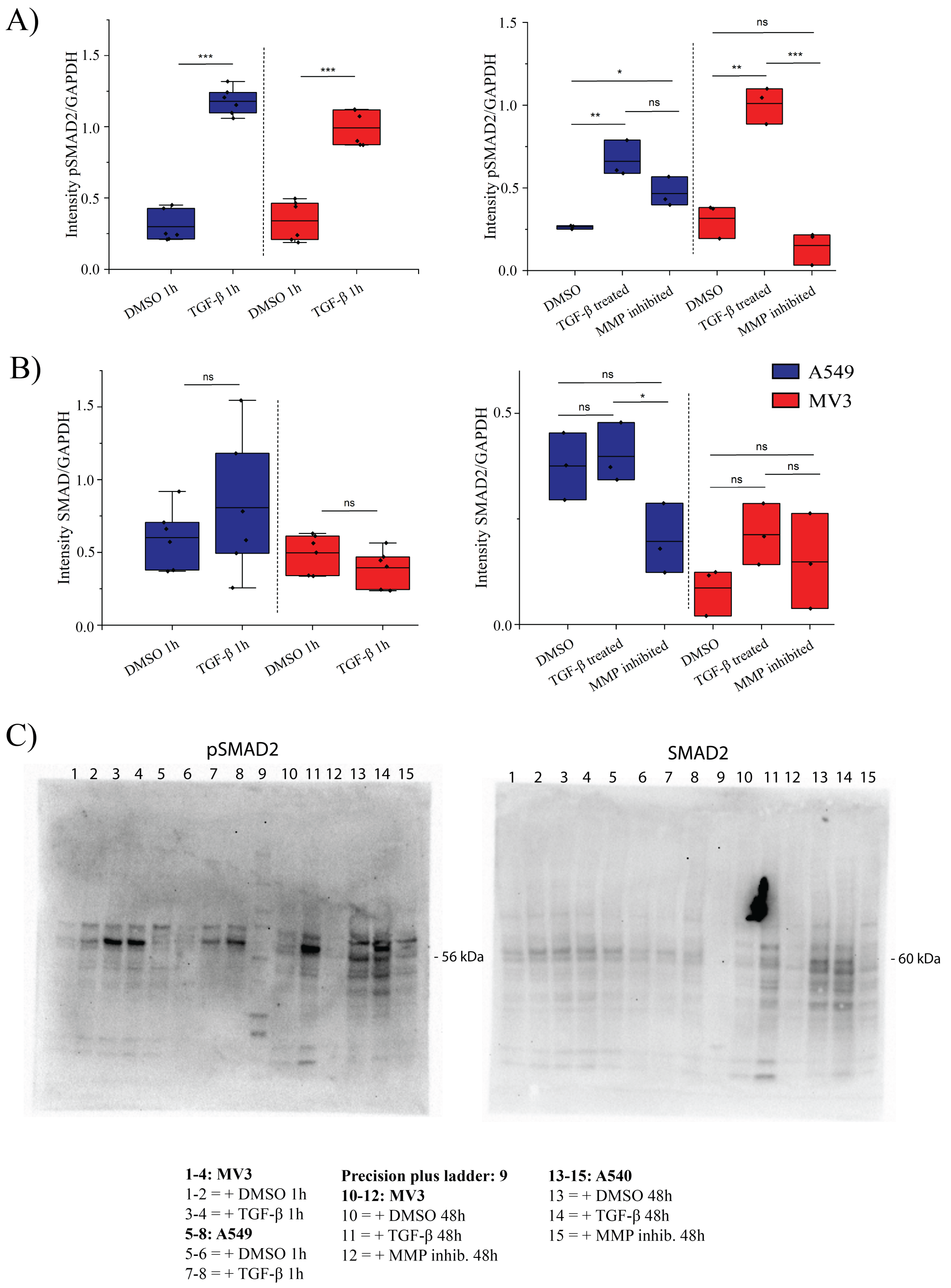




Figure 13: **Western Blot analysis for SMAD2 and pSMAD2 expression of MV3 and A549 cells cultured in 2D on collagen coated 6-well plates.** A) Protein expression of pSMAD2 after 1h (left) and 48h (right) of TGF-$\beta$ treatment in A549 and MV3 cells. B) Protein expression of SMAD after 1h (left) and 48h (right) of TGF-$\beta$. C) Western blot images of pSMAD2 (left) and SMAD2 (right). The legend shows the content of each lane (numbered on top of the gels). For 1h treatments: (n=2). For 48h treatments: (n=1).

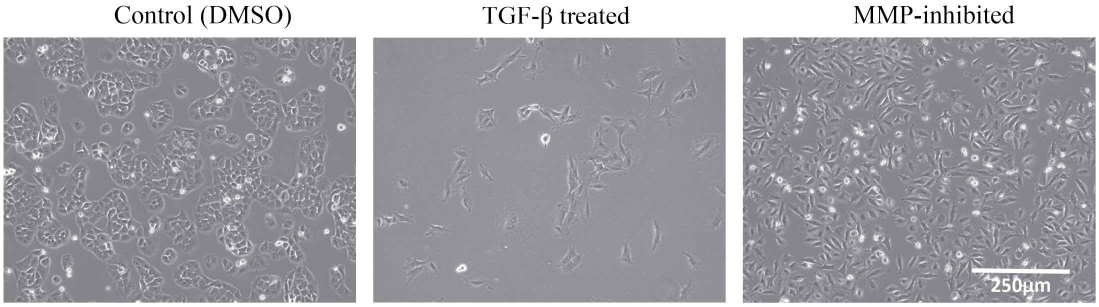


Figure 14: **A549 cell morphology in 2D cell culture under control (DMSO), TGF-$\beta$ treated and MMP-inhibited conditions.** Bright field images were taken 2 days after seeding on collagen coated 6-well plates. Scale bar: 250 $\mu$m.

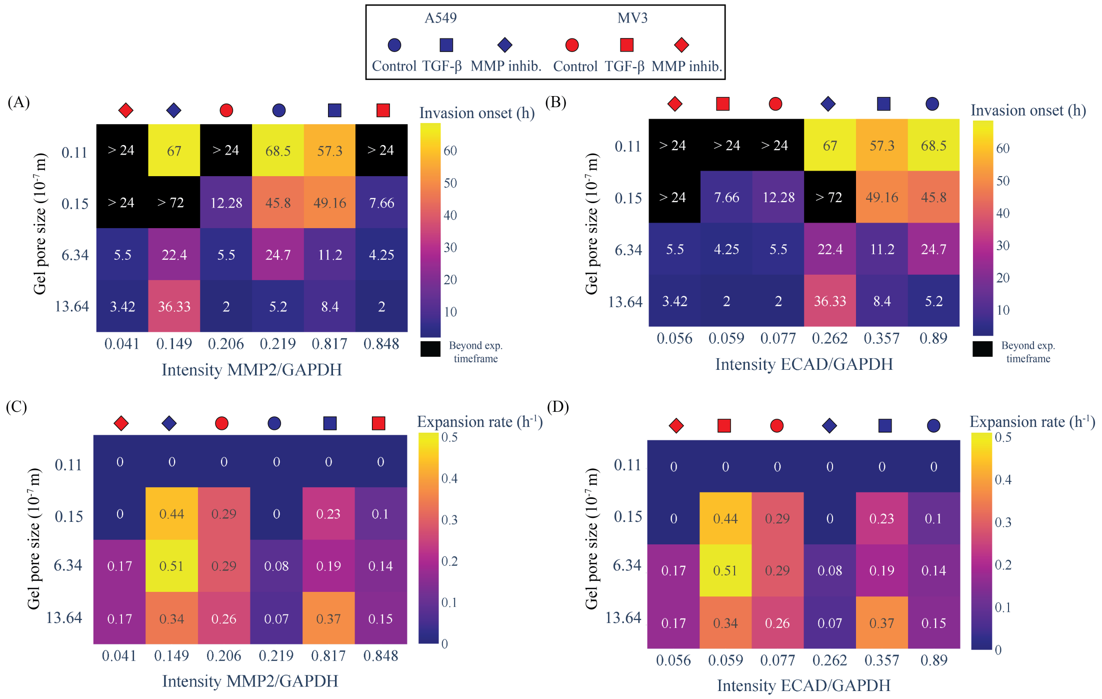


Figure 15: **Correlation of the invasion onset times and spheroid expansion rates with MMP2 and E-cadherin protein expression levels for MV3 and A549 spheroids.** A) Heat map showing the dependence of the invasion onset time (indicated by the number (in units of hrs) in each square and the color code, see color bar on the right) on hydrogel pore size (y-axis) and expression level of MMP2 (x-axis). B) Corresponding heat map showing the dependence of the invasion onset time on hydrogel pore size (y-axis) and expression level of E-cadherin (x-axis). Note that spheroids that did not invade during the time frame of the assay are shown in black. C) Heat map showing the dependence of the spheroid expansion rate (number (in units of $hr^{-1}$) in each square and color code, see color bar on the right) on hydrogel pore size (y-axis) and expression level of MMP2 (x-axis). D) Corresponding heat map showing the dependence of the spheroid expansion rate on hydrogel pore size (y-axis) and expression level of E-cadherin (x-axis). In all cases, data for MV3 and A549 spheroids were pooled and protein expression levels were normalized by GAPDH.

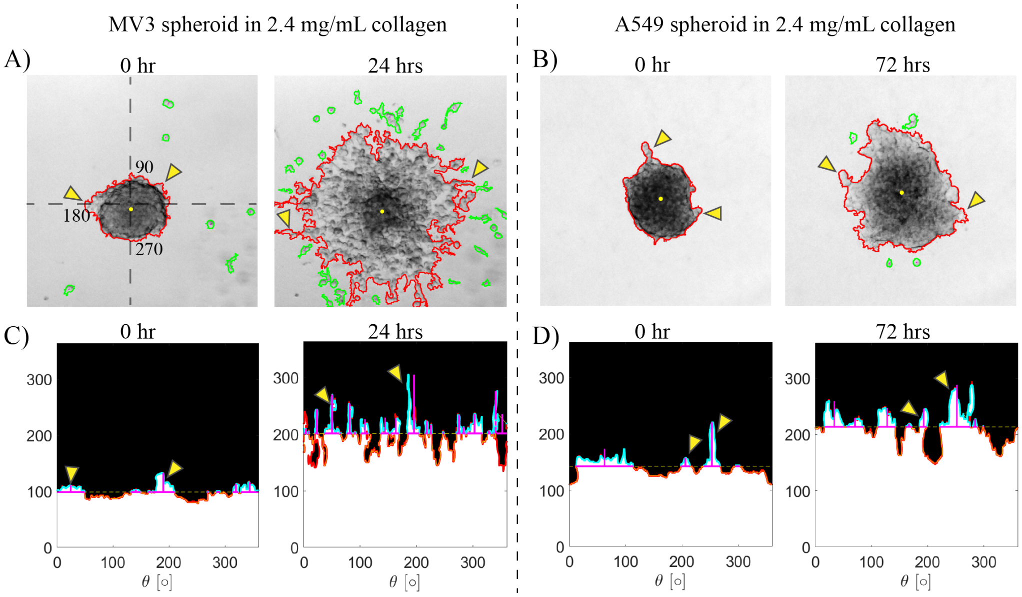


Figure 16: **Analysis of spheroid protrusions and single cell dissemination aimed at classifying the state (gas, fluid, liquid) of spheroids under different conditions.** A) Bright field images of a MV3 spheroid in 2.4 mg/mL collagen at t = 0 (initial state) and t=24 hrs (final state). B) Bright field images of an A549 spheroids in 2.4 mg/mL collagen at t=0 hr (initial state) and t=72 hrs (final state). Spheroid boundaries are outlined in red while disseminated cells are outlined in green. The yellow arrowheads highlight examples of multicellular protrusions that grow over time. Note that both MV3 and A549 cells already showed some protrusions at t=0 hr. Additionally, the images often revealed some disseminated cells at t=0 hr. Therefore, in all cell count quantification, we subtracted the cells present at t=0 hr. C) Polar plots determined from the MV3 spheroid images in A, based on the assumption of radial symmetry. D) Corresponding polar plots for the A549 spheroid images in B. In both cases, multicellular protrusions are seen as peaks (yellow arrowheads highlight examples). The polar plots were used as a basis to detect each peak (i.e., multicellular protrusion) and quantify its maximum height relative to the average spheroid radius (horizontal red lines). Spheroids with average protrusion lengths greater than 25 $\mu$m were considered to be in a liquid-like (unjammed) state.

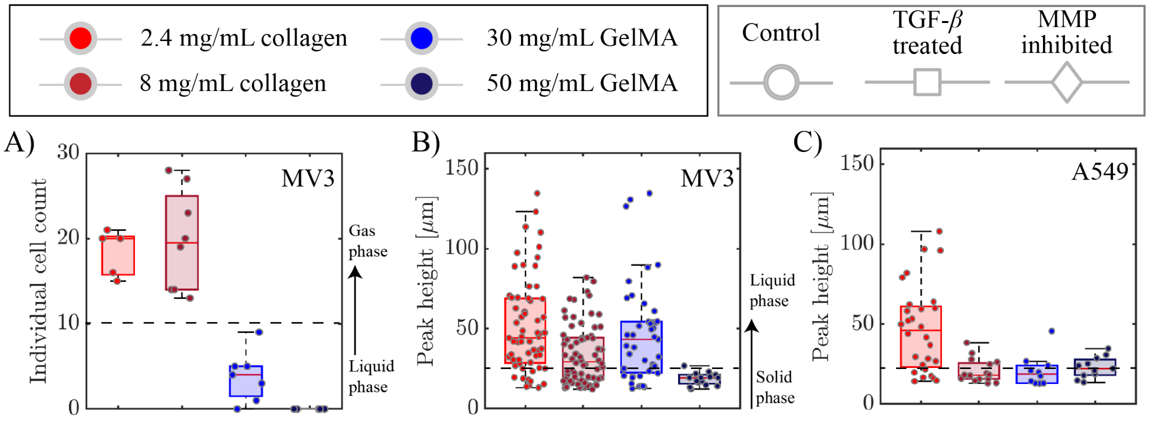


Figure 17: **Analysis of spheroid protrusions and single cell dissemination aimed at classifying the state (gas, fluid, liquid) of MV3 and A549 spheroids in control conditions.** A) Individual cell count analysis of MV3 spheroids in collagen (2.4 and 8 mg/mL) and in GelMA (30 and 50 mg/mL) matrices. Spheroids are considered to be in a gas-like state when the cell count is above 10 (above the horizontal dashed line) and solid-like or liquid-like when the cell count is below 10. MV3 spheroids were gas-like in collagen (2.4 and 8 mg/mL) and solid-like in GelMA (30 and 50 mg/mL GelMA). B) Protrusion lengths for MV3 spheroids. C) Protrusion lengths for A549 spheroids. Spheroids are considered liquid-like when the protrusion lengths are above 25 $\mu$m (above the dashed lines) and solid-like when the protrusion lengths are below 25 $\mu$m. MV3 spheroids were gas-like in collagen (2.4 and 8 mg/mL) and 30 mg/mL GelMA hydrogels and solid-like in 50 mg/mL GelMA. A549 spheroids were liquid-like only in 2.4 mg/mL collagen and solid-like otherwise.

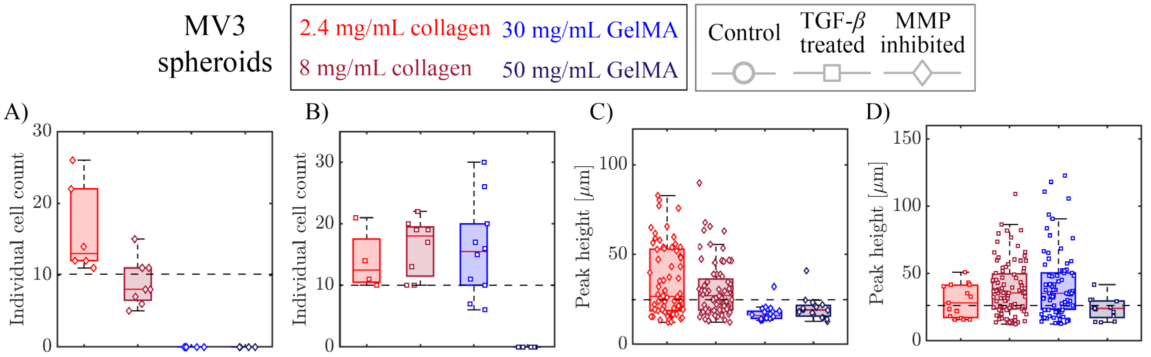


Figure 18: **Quantification of disseminated cells and protrusion lengths for MV3 spheroids in order to classify the state (gas, fluid, solid) of the spheroid.** A) Individual cell count analysis for MV3 spheroids treated with MMP-inhibitor in different collagen and GelMA gels (see legend on top). MMP-inhibitor treatment caused a strong (40-50%) drop in the cell count in collagen (2.4 mg/mL and 8 mg/mL) gels and blocked cell dissociation in 30 GelMA hydrogels. B) Corresponding data for MV3 cells treated with TGF-$\beta$. The dashed lines indicate the transition between a liquid/solid state (cell count below 10) and a gas state (cell count above 10). TGF-$\beta$ treatment caused a large (73%) increase in cell count in 30 mg/mL GelMA compared to control conditions but did not result in any individual cell dissemination in 50 mg/mL GelMA. C) Protrusion lengths for MV3 spheroids treated with MMP-inhibitor in different collagen and GelMA gels. D) Corresponding data for MV3 spheroids treated with TGF-$\beta$. The dashed lines indicate the transition between a solid state (maximum protrusion length below 25 $\mu$m) and a liquid state (average protrusion length above 25 $\mu$m). Box plots show N = 5-10 spheroids per condition performed in 2 independent experiments.

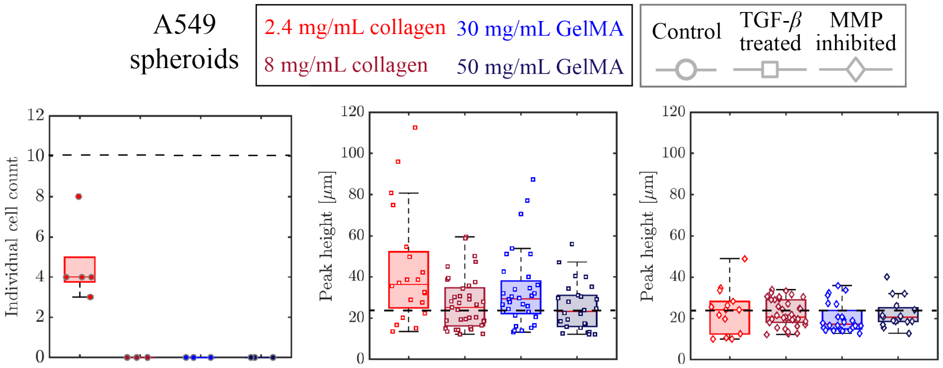


Figure 19: **Quantification of disseminated cells and protrusion lengths for A549 spheroids in order to classify the state (gas, fluid, solid) of the spheroid.** A) Individual cell count analysis for A549 spheroids embedded in collagen and GelMA matrices (see legend on top) in control conditions. The dashed line indicate the transition between a liquid/solid state (cell count below 10) and a gas state (cell count above 10). Under control conditions, the A549 spheroids only disseminated individual cells in 2.4 mg/mL collagen. Note that A549 spheroids embedded in 2.4 mg/mL collagen treated with TGF-$\beta$ disintegrated and sedimented to the bottom, so we were unable to count the disseminated cells. At early times, however, we could observe cell dissociation, so we classify this condition as a gas-like phase. B) Protrusion lengths of A549 spheroids treated with TGF-$\beta$. A549 spheroids formed protrusions characteristic of a liquid-like state in 8 mg/mL collagen and in 30 mg/mL GelMA upon TGF-$\beta$ treatment, whereas they were solid-like in control conditions. In 8 mg/mL collagen, the TGF-$\beta$-treated spheroids made shorter protrusions with an average length of 26 $\mu$m. No protrusions were observed for spheroids embedded in 50 mg/mL GelMA. C) Protrusion lengths of A549 spheroids treated with MMP-inhibitor. Upon MMP inhibition, A549 spheroids were liquid-like only in 2.4 mg/mL collagen. The dashed lines in B and C indicate the transition between a solid state (average protrusion length below 25 $\mu$m) and a liquid state (average protrusion length above 25 $\mu$m). Box plots show N = 5-10 spheroids per condition.

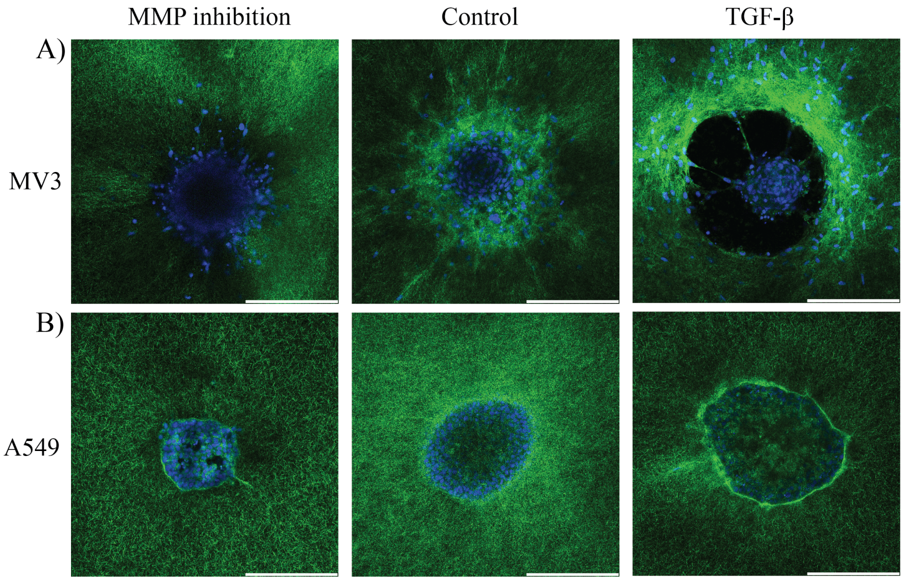


Figure 20: **Visualization of cell-mediated extracellular matrix remodelling for spheroids embedded in collagen (2.4 mg/mL).** Confocal images of A) MV3 and B) A549 spheroids in control conditions (middle) and with MMP-inhibition (left) or TGF-$\beta$ treatment (right) around spheroid equator. Collagen fibers were imaged by reflection and are shown in green. Nuclei stained with Hoechst were imaged by fluorescence and are shown in blue. Scale bars are 250 $\mu$m. Compared to control conditions, TGF-$\beta$ stimulation causes more matrix remodeling whereas MMP inhibition diminishes matrix remodeling. MV3 spheroids appear to exert higher traction forces on the collagen than A549 spheroids, as shown by radially oriented fibers around the spheroid under MMP inhibition and control conditions and extensive collagen accumulation upon TGF-$\beta$ stimulation. Furthermore, the black void around the MV3 spheroid upon TGF-$\beta$ stimulation is indicative of MMP-mediated matrix degradation.

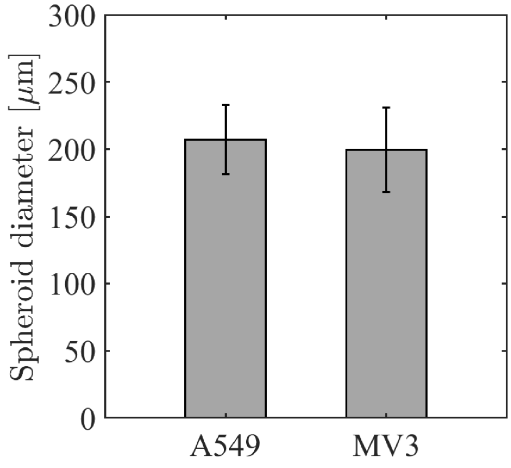


Figure 21: **Average diameter of MV3 (N = 80) and A549 (N = 85) spheroids at t=0 hr determined from bright field images.** The spheroids had comparable average diameters of 200±30 $\mu$m (MV3) and 207±26 $\mu$m (A549).
